# Inflammatory Neuropathy in Mouse and Primate Models of Colorectal Cancer

**DOI:** 10.1101/2024.07.19.604274

**Authors:** Caitlyn M. Gaffney, Angela M. Casaril, Iqbal Mahmud, Bo Wei, Karen M. Valadez, Elizabeth A. Kolb, Fisher R. Cherry, Theresa A. Guise, Philip L. Lorenzi, Lei Shi, Carolyn L. Hodo, Andrew J. Shepherd

## Abstract

Colorectal cancer survivors are at increased risk of developing neurological issues, particularly peripheral neuropathy and chronic pain. Although pre-existing neuropathy is a risk factor for chronic pain, tumor-induced neuropathy has not been firmly established in pre-clinical models. Consistent with clinical observations, we show that mice with colorectal cancer develop peripheral neuropathy, which was associated with subtle locomotor deficits, without overt hypersensitivity. We detected widespread differences in pro-inflammatory cytokines and lipid metabolites in peripheral nerves from tumor-bearing mice. Macrophage accumulation, myelin decompaction and ryanodine receptor oxidation were associated with dysfunctional calcium homeostasis and reduced spike amplitude in sensory neurons. Inflammatory neuropathy and macrophage accumulation were also observed in peripheral nerves of rhesus macaques with colorectal cancer. These findings suggest colorectal cancer is causally linked to a subacute form of chronic inflammatory demyelinating polyneuropathy across species, which may represent an under-reported, yet important risk factor for neurological dysfunction in colorectal cancer survivors.

## Introduction

There were an estimated 150,000 diagnoses of colorectal cancer (CRC) in the US in 2023^1^. Advances in screening and treatment have improved CRC survival, with 5-year survival for localized disease above 90% and more than 1.4 million total CRC survivors in the US in 2022^2^. This success inevitably prompts greater focus on long-term complications of cancer and its treatment, especially neuropathy and pain. Overall pain prevalence in cancer survivors is estimated at 45-60%^3^, with survivors frequently reporting widespread, persistent pain for months to years after completing treatment. The major precipitating factors in this pain are invasive surgery and chemotherapy-induced peripheral neuropathy^4^. In addition to a lack of effective pharmacotherapies for neuropathic pain in cancer survivors^5^, there is also a dearth of understanding surrounding the factors that increase the risk of developing chronic pain. However, prior nervous system injury is one such risk factor^6^. This injury has many potential causes, including surgery, trauma, metabolic dysfunction and - crucially - cancer itself. Cancer-related or paraneoplastic neuropathy is clinically defined by neurological symptoms due to autoantibody production against neuronal antigens and is a well-established, albeit rare sequela of lymphoma, lung, and breast cancer^7^. However, the extent to which other neoplasms drive neuropathy is largely unknown, and the widespread underdiagnosis of many forms of neuropathy^8^ raises the likelihood that paraneoplastic neuropathy is similarly under-reported. Additionally, two prior studies of CRC patients found somewhat unexpected sensory deficits and reduced cutaneous nerve fiber density before beginning chemotherapy^9, 10^, hinting that neuropathy may be a common and therefore under-diagnosed complication of CRC. Determining the mechanistic basis of this neuropathy is an important step toward accurately assessing the risk of neurological complications and other adverse events due to CRC and its treatment.

Our previous work showed reduced cutaneous nerve fiber density and sensory neuron mitochondrial dysfunction in a mouse model of CRC, both hallmarks of peripheral neuropathy^11^. We also detected increased pro-inflammatory cytokines in serum, but the behavioral effects of this neuropathy and the implications for peripheral nerve inflammation are not fully understood. We now show that an immunocompetent, orthotopic model of microsatellite instability (MSI-H) CRC exhibits neuropathy which manifests in impaired motor coordination, dyslipidemia, inflammation and demyelination, along with dysfunctional Ca^2+^ homeostasis in sensory neurons. We also recapitulate the neuroinflammatory, neurodegenerative and demyelinating effects of CRC in peripheral nerve biopsies from rhesus macaques with spontaneous MSI-H CRC. Because pre-existing neuropathy driven by CRC may represent a major risk factor for subsequent neurological issues, these findings have hitherto-unrecognized implications for quality-of-life in cancer survivorship.

## Results

### MC38 tumor mice show no alterations in tactile and thermal sensitivity, with deficits in fine motor skills

Orthotopic MC38-luciferase tumor burden (as measured by luciferase signal) did not significantly differ between sexes (Fig. 1a-b), and significant loss of PGP9.5-immunoreactive intraepidermal nerve fibers (IENFs) was detected in hindpaws of female MC38 tumor mice versus vehicle controls (*t*(26) = 2.383, *p* = 0.025; Fig. 1c-d), consistent with our prior observations in males^11^. Tumor growth was not associated with significant differences in von Frey hindpaw withdrawal threshold (Fig. 1e). Tumor burden and paw withdrawal threshold did not significantly correlate (Fig. 1f), but there was a trend toward reduced withdrawal threshold with increasing tumor burden specifically in males (R^2^ = 0.12, F_1,31_ = 4.025, *p* = 0.054; Supplementary Fig. 1a). The opioid antagonist naltrexone was used to unmask any potential latent pain sensitization^12^ in MC38 mice. There was a significant, albeit brief effect of tumor/naltrexone dosing (F_3,18_ = 3.472, *p* = 0.038; Fig. 1g). Specifically, paw withdrawal threshold was lower 15 minutes post-naltrexone in MC38 mice versus controls (*p* = 0.014; Tukey’s multiple comparisons test), but this effect was also present in pre-naltrexone baseline readings (*p* = 0.035; Tukey’s multiple comparisons test), and was absent at 30-, 90-, and 120-minutes post-dosing (Fig. 1g). Performance in the burrowing assay, an activity that is suppressed by pain^13^, was also comparable between MC38 mice and vehicle controls (Fig. 1h).

**Figure 1.**
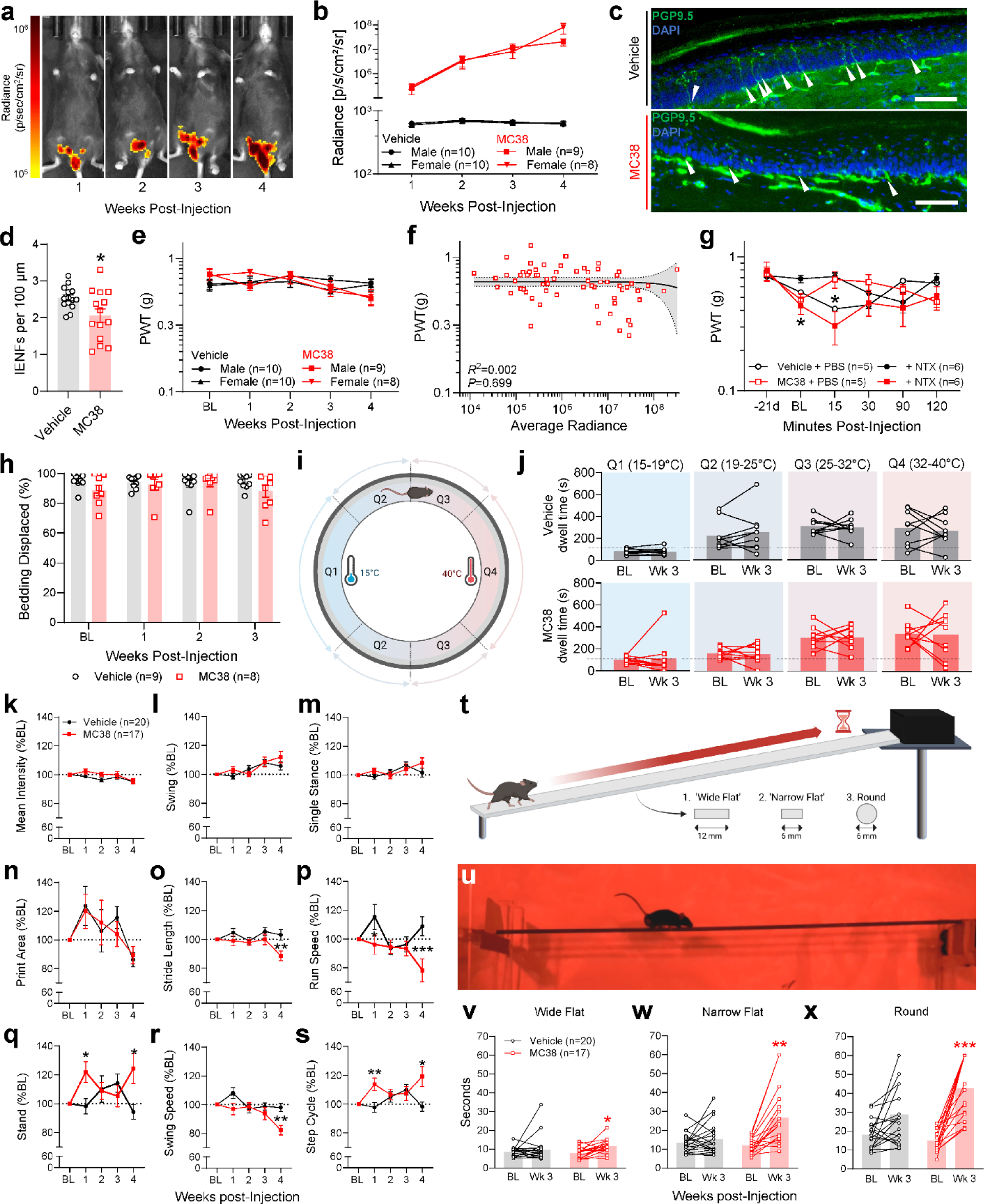
MC38 tumor mice show no clear changes in tactile and thermal sensitivity, with deficits in fine motor skills. (**a**) Bioluminescent imaging shows increasing colorectal tumor burden up to 4 weeks after orthotopic injection of MC38 cells. (**b**) Quantification of luciferase signal intensity shows MC38 tumor growth is approximately equal in male and female mice. (**c**) Representative images and quantification (**d**) show significantly reduced density of PGP9.5-immunoreactive IENFs (white arrowheads) in MC38 mouse hindpaw skin at week 3 post-injection. Error bars = SEM, * = *p* <0.05, vehicle vs MC38, unpaired t-test, scale bar = 100 µm. (**e**) No significant differences in paw withdrawal threshold (PWT) in male or female mice bearing MC38 tumors. (**f**) No correlation between MC38 luciferase signal and paw withdrawal threshold across time. (**g**) Three weeks post-vehicle or MC38 injection, mice received subcutaneous vehicle (PBS) or naltrexone (NTX, 3 mg/kg) to unmask any latent pain hypersensitivity. In MC38-injected mice, NTX was associated with a significant reduction in PWT versus PBS-injected controls at 15 minutes post-injection only. Error bars = SEM, * = *p* <0.05, ‘MC38 + NTX’ versus ‘Vehicle + NTX’. Two-way repeated measures ANOVA, Tukey’s multiple comparison test. (**h**) Burrowing activity does not significantly differ between vehicle-injected and MC38-injected mice. (**i-j**) Thermal gradient ring assay measures temperature preference between 15 and 40°C. Vehicle and MC38-injected mice show no significant differences in dwell time from baseline to 3 weeks post-injection, with similar aversion to Q1 (15-19°C) and preference for Q3 and Q4 (25-32°C and 32-40°C, respectively). (**k-s**) Voluntary gait analysis revealed no significant differences between vehicle and MC38-injected mice in hindpaw (**k**) mean print intensity, (**l**) swing time (time lifted off the ground), (**m**) single stance time (time one hindlimb spends bearing weight) or (**n**) print area. Predominantly late-stage deficits were detected in (**o**) stride length, (**p**) average run speed, (**q**) stand time, (**r**) swing speed and (**s**) step cycle time. Error bars = SEM. (**t**) illustration of the beam walk task and cross-sectional profile of the three increasingly challenging beams. (**u**) video still of a mouse traversing the round beam. (**v**) Three weeks post-vehicle injection, there was no significant difference versus baseline in traversal time of the wide flat beam. However, significant increases in crossing time were seen in MC38-injected mice with the wide flat, (**w**) narrow flat and (**x**) round beams. */**/*** *p* = <0.05 / 0.01 / 0.001 ‘week 3’ versus baseline ‘BL.’ Paired T-test, Bonferroni correction.

Neuropathic and inflammatory pain states commonly feature cold and heat hypersensitivity, respectively^14^. Using a 15-40°C thermal gradient ring^15^ (Fig. 1i), we did not detect any shift in temperature preference or aversion in weeks 1 and 2 post-MC38 injection compared to baseline (Supplementary Fig. 1b). This was also true 3 weeks post-injection (Fig. 1j), a timepoint at which IENF loss is evident.

Peripheral neuropathy is also an independent risk factor for gait abnormalities and impaired balance^16^. We saw that pawprint intensity, swing time, hindpaw single stance time and print area did not significantly differ between vehicle and MC38 mice (Fig. 1k-n). However, there was an effect of tumor growth on stride length (F_1,38_ = 4.400, *p* = 0.043; Fig. 1o) which was significant at 4 weeks (*p* = 0.013, mixed effects analysis). Tumor growth also reduced run speed (F_1,38_ = 4.234, *p* = 0.047) in weeks 1 and 4 (*p =* 0.004, *p* = < 0.001; Fig. 1p). Stand time was increased in tumor-bearing mice (F_1,38_ = 2.508, *p* = 0.0122), in weeks 1 and 4 (*p* = 0.004, *p* = 0.012; Fig. 1q). Conversely, swing time was reduced in tumor-bearing mice (F_1,38_ = 7.074, *p* = 0.011) in weeks 1 and 4 (*p* = 0.018, *p* = 0.001; Fig. 1r). Since overall step cycle time is the sum of stand and swing time, this measure was also impacted by tumor growth (F_1,38_ = 5.323, *p* = 0.027) and was significantly greater in tumor-bearing mice in weeks 1 and 4 (*p* = 0.003, *p* = 0.007; Fig. 1s). Except for stand index (F_1,38_ = 1.996, *p* = 0.166; week 4: *p* = 0.009) and body speed (F_1,38_ = 6.164, *p* = 0.018; week 1: *p* = 0.014; week 4: *p* = < 0.001), other gait measures were not significantly affected (Supplementary Fig. 1c). We detected main effects of sex and presence of MC38 tumor in run speed (F_1,36_ = 8.894, *p* = 0.005 [sex], 6.081, *p* = 0.019 [tumor]) and step cycle time (F_1,36_ = 4.921, *p* = 0.033 [sex], 5.444, *p* = 0.025 [tumor]). Sex differences between MC38-injected mice were seen in swing speed (F_3,36_ = 3.090, *p* = 0.039), which was lower in females at week 3 (p = 0.041). Otherwise, there were no significant, sustained differences between male and female MC38 mice in development of these deficits (Supplementary Fig. 1d).

Because gait analysis measures normal, voluntary locomotor function, we subjected mice to the more demanding beam walk task, which measures traversal time of three increasingly challenging beams (Fig. 1t-u). Three weeks post-injection, MC38 mice took significantly longer to cross all three beams compared to their pre-injection baselines (‘wide flat’: *t*(16) = 3.141, *p* = 0.038; *‘*narrow flat’: *t*(16) = 4.647, *p* = 0.002; ‘round’: *t*(16) = 6.289, *p* = 0.0006, Bonferroni correction; Fig. 1v-x and Supplementary Video Files 1-6). There were no clear sex differences in beam walk performance (Supplementary Fig. 1e). Although mice with the greatest performance decrement tended to have MC38 tumors, the average number of errors (slips, inversions, falls) did not significantly increase post-MC38 injection (Supplementary Fig. 1f). This suggests MC38 tumor-bearing mice exhibit histological signs of peripheral neuropathy in skin, without overt or latent hypersensitivity to tactile or thermal stimuli, but with subtle impairment of motor coordination.

### Hypercoagulability and dyslipidemia in MC38 tumor mice

We next determined whether MC38 tumor mice develop biomarkers of hyperglycemia, hypercoagulability, or renal/hepatic failure, since they are all potential complications of cancer as well as neuropathy^17^. We observed no tumor-related differences in blood glucose, but there was a trend toward reduced bleeding time and blood loss in tumor-bearing mice, which was accompanied by significantly increased fibrinogen and a trend toward reduced fibrinogen time (Supplementary Fig. 2a). Other coagulation measures (prothrombin time, antithrombin III, activated partial thromboplastin time [aPTT]) did not significantly differ (Supplementary Fig. 2b).

With the exception of aspartate transaminase (*t*(6) = 2.470, *p* = 0.049), markers of hepatic function (alkaline phosphatase, gamma-glutamyl transferase, alanine transaminase; Supplementary Fig. 2c) and renal function (phosphorus, sodium, potassium, creatinine, albumin, total protein; Supplementary Fig. 2d) were not altered by MC38 tumor growth. This suggests that although liver and kidney failure can lead to neuropathy, neither are likely causes of MC38-related neuropathy. Finally, lactate dehydrogenase (LDH), but not creatine kinase (CK) showed a strong trend toward elevation in MC38 mice (LDH: *t*(6) = 2.395, *p* = 0.054; Supplementary Fig. 2e). LDH and CK are markers of CRC burden and muscle degeneration, respectively^18, 19^.

CRC and MC38 tumor growth both cause systemic inflammation^11^, which is linked to dyslipidemia. We next established whether dyslipidemia could underlie the neuropathy caused by MC38 tumor growth. Of the 371 lipid species detected in MC38 mouse plasma, 13 were significantly upregulated and 101 were significantly downregulated versus controls (Fig. 2a and Supplementary Table 1). MC38 mouse plasma showed significant and widespread triglyceride depletion (TG; Fig. 2b-c). Long-chain fatty acids (FAs; >18 carbons) and unsaturated FAs were also enriched in MC38 mouse plasma, while comparatively shorter chain lengths (13-18 carbons) and unsaturated FAs were reduced (Fig. 2d-e). Numerous ‘inflammatory lipid’ families (sphingolipids, sphingomyelins, ceramides, hexosylceramides, cholesterol esters) were enriched in MC38 mouse plasma, along with organelle-specific lipid ontologies (‘mitochondrion’, ‘endoplasmic reticulum’, ‘endosome/lysosome’; Fig. 2f-h). Around two-thirds of the phosphatidylcholine (PC) species detected were enriched in MC38 mouse plasma, whereas two-thirds of phosphatidylethanolamines (PE) were reduced (Fig. 2i-j). As expected, enrichment and reduction in PC and PE was reflected in the levels of their products: lysophosphatidylcholine (LPC) and lysophosphatidylethanolamines (LPE; Fig. 2k). Finally, phosphatidylinositol (PI) species were both enriched and reduced in tumor-bearing mice, with no clear distinction in chain length or saturation index (Fig. 2l). Broadly similar changes in plasma lipid enrichment and depletion were replicated in an independent cohort of vehicle and MC38 tumor-bearing mice (Supplementary Fig. 3).

**Figure 2.**
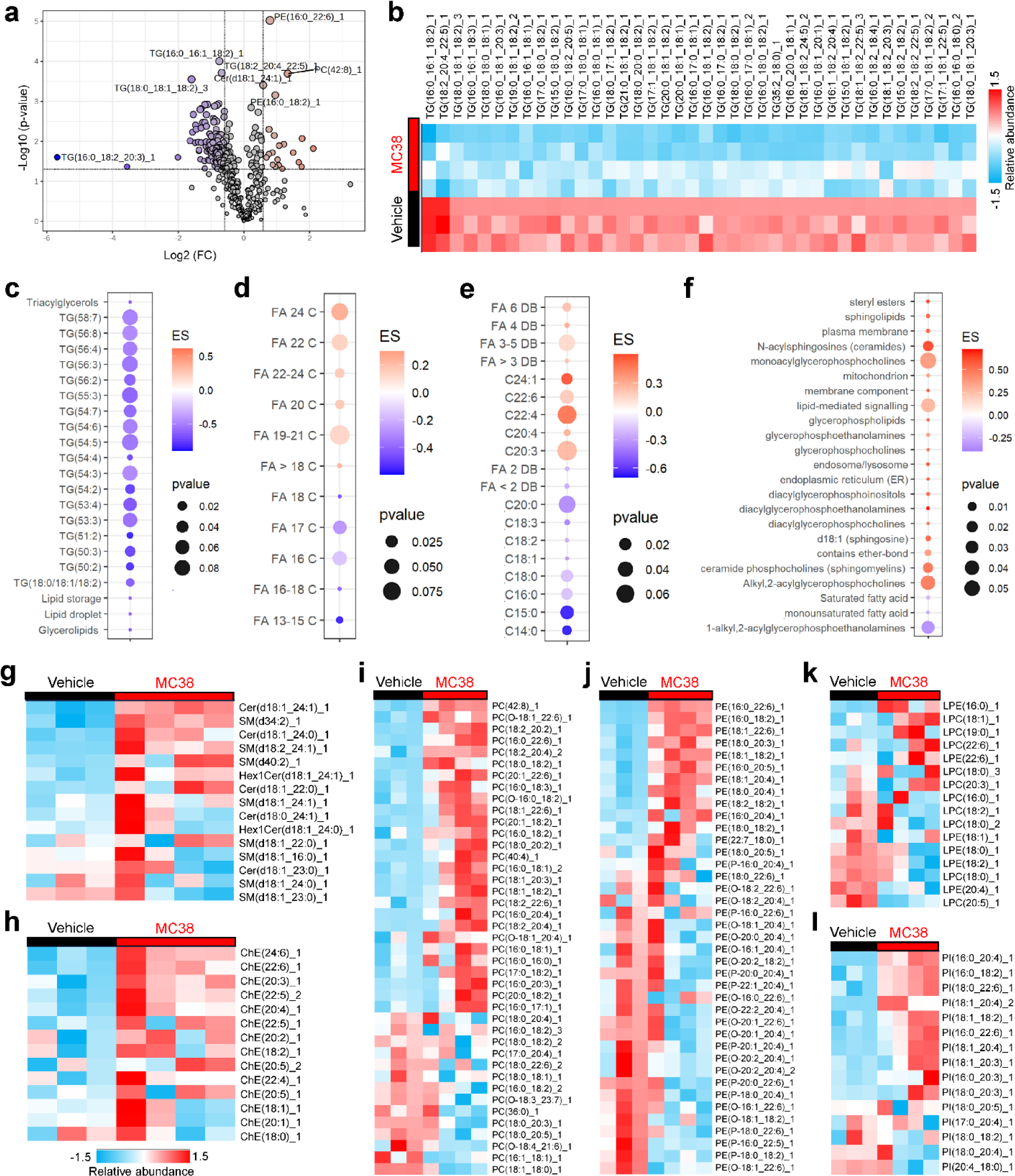
MC38 tumor growth is associated with dyslipidemia. (**a**) Volcano plot displays log2 fold-change (threshold ≥1.5) and p-value (threshold ≤0.05) of lipid abundance in MC38 tumor-bearing mouse plasma versus control. (**b**) Heatmap and enrichment score (ES; **c**) shows widespread reduction of triglycerides in tumor-bearing mice. (**d**) Chain length analysis shows an enrichment of long-chain fatty acids and reduction of shorter-chain molecules. (**e**) Saturation index shows enrichment of several polyunsaturated fatty acids alongside reduction of saturated and monounsaturated fatty acids. (**f-g**) Analysis of lipid class enrichment reveals increased ceramides and sphingomyelins. (**h**) Cholesterol esters are enriched in plasma from tumor-bearing mice (**i**) Phosphatidylcholines (PC) are majority enriched in MC38 plasma (**j**) Phosphatidylethanolamines (PE) are majority reduction in MC38 plasma. (**k**) Lysophosphatidylcholine/ethanolamine show similar enrichment/depletion dynamics to PC/PE (**l**) Phosphatidylinositol (PI) abundance is highly variable, with some enrichment and some depletion in MC38 tumor-bearing mice.

### Alterations in lipid composition consistent with neuropathy in nerves from tumor-bearing mice

The profound dyslipidemia in MC38 mouse plasma is consistent with prior reports of malignancy-associated dyslipidemia in mice and humans^20, 21^. We next surveyed the lipid composition of the sciatic nerve, which is known to be disrupted by plasma dyslipidemia in demyelinating diseases^22^. Consistent with results in plasma, we observed 250 enriched and 50 reduced lipids in sciatic nerves from MC38 mice, along with a significant reduction in TGs (Fig. 3a-c and Supplementary Table 2). Longer-chain FA enrichment and medium-length FA depletion seen in plasma was mirrored in nerve tissue, as was enrichment of mitochondria and endoplasmic reticulum-related lipid species (Fig. 3d-e). Inflammatory lipids including sphingolipids, sphingomyelin and ceramides were also significantly enriched (Fig. 3f-g). As with plasma lipids, an independent experiment reproduced these changes in nerve lipid composition (Supplementary Fig. 4).

**Figure 3.**
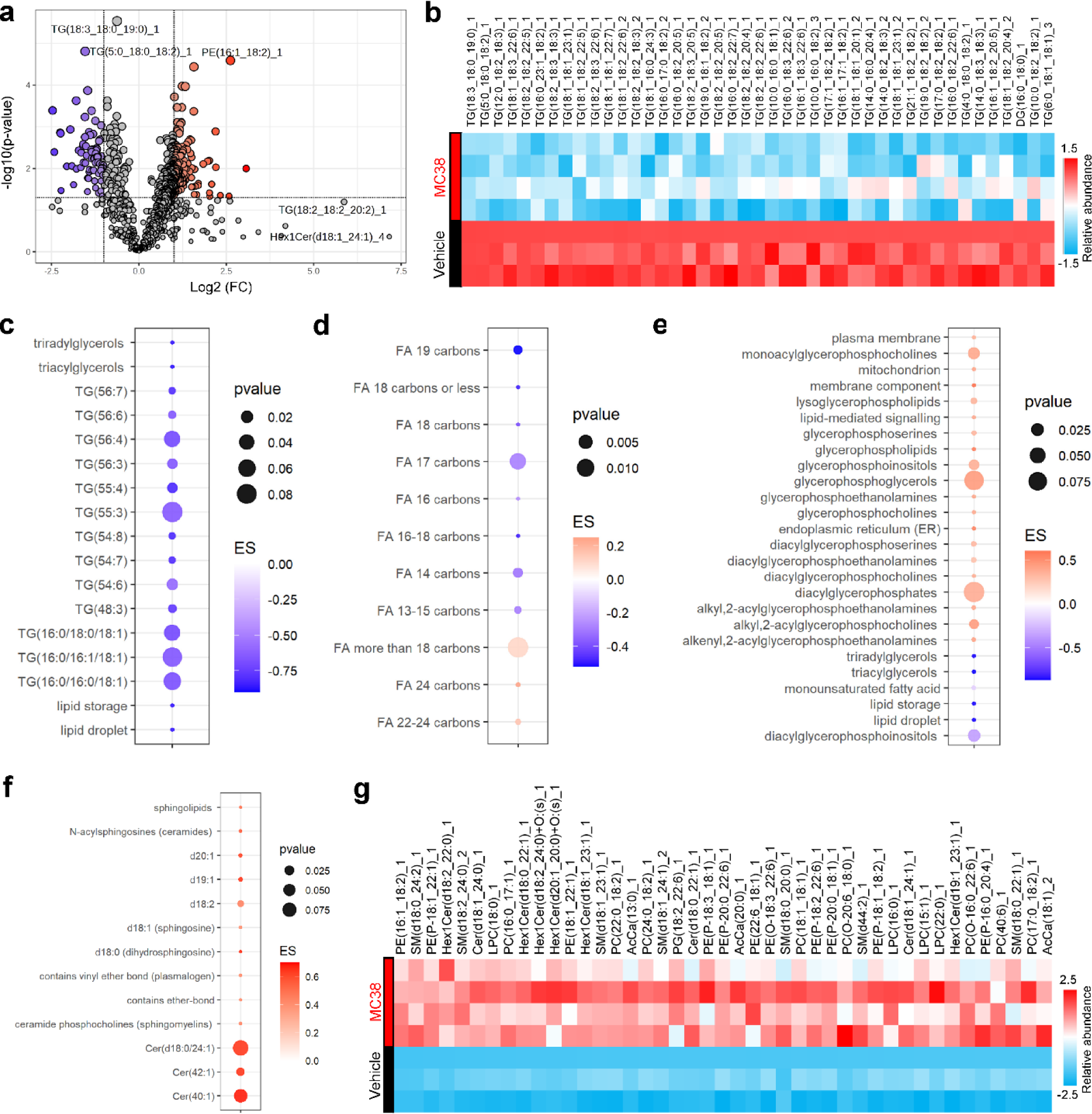
Lipidomic analysis of sciatic nerves from MC38 tumor-bearing mice. (**a**) Volcano plot displaying log2 fold-change (threshold ≥1.5) and p-value (threshold ≤0.05) of lipid abundance in MC38 tumor-bearing mouse sciatic nerve versus control. (**b-c**) Heatmap and enrichment scores show widespread reduction of triglycerides in tumor-bearing mice. (**d**) Chain length analysis shows an enrichment of long-chain fatty acids and reduction of shorter-chain molecules. (**e**) Analysis of lipid class enrichment reveals increased diacylglycerolipids and reduced triacylglycerols. (**f**) Sphingolipids and ceramides are significantly enriched in MC38 nerves. (**g**) Phosphatidylethanolamines (PE), sphingomyelin, PC, ceramides and hexosylceramides are enriched in sciatic nerves from MC38 tumor bearing mice compared to control mice.

Importantly, increased gut permeability is associated with inflammation and dyslipidemia^23^. To test whether CRC and associated damage to the gastrointestinal tract was causing systemic inflammation and dyslipidemia, we orally dosed mice with a fluorescent dextran conjugate. No increase in fluorescence was detected in the blood of MC38 mice versus controls 4h later, consistent with gastrointestinal integrity being broadly intact in MC38 mice. Additionally, a luminol test did not detect occult fecal bleeding (Supplementary Fig. 5a-b), suggesting that frank anemia is also unlikely to underlie MC38-driven neuropathy. Histologically, there was some evidence of increased cellularity in colonic submucosa (Supplementary Fig. 5c-g), but no obvious loss of integrity or perforation. This suggests that MC38-driven neuropathy and dyslipidemia are associated with symptoms of gastrointestinal inflammation, but no overt loss of barrier integrity.

### Evidence of Schwann cell injury and inflammation in MC38 mouse sciatic nerves

The substantial peripheral nerve lipid disruptions prompted us to re-evaluate pre-existing bulk sequencing datasets of sensory ganglia from MC38 tumor-bearing mice at the same timepoint. Consistent with the lipidomic findings, there was significant differential expression of genes related to lipid metabolism and Schwann cell function (e.g. *Car3, Lpl, Mag, Mpzl2, Myrf, Plin4, Pltp, Pmp22, Smpd2*; Supplementary Fig. 6a)^24, 25^. Collectively, this suggests that MC38 tumor growth either directly or indirectly distorts lipid metabolism in ways that may lead to neuronal/Schwann cell dysfunction.

Because we detected lipidomic and transcriptional changes that suggest Schwann cell dysfunction, we stained sciatic nerve sections for S100B, which is secreted by activated Schwann cells in neurodegeneration^26^. S100B immunoreactivity was elevated in sciatic nerves from MC38 mice (*t*(22) = 2.701, *p* = 0.013; Fig. 4a-c). Along with the profound dysregulation of neuronal lipid composition, this led us to examine nerve ultrastructure using electron microscopy. Vehicle controls showed electron-dense, compact myelin around large caliber axons (Fig. 4d-e). However, a significant increase in myelin decompaction (‘onion bulb’ formations) was seen in MC38 mouse nerves (*t*(22) = 5.620, *p* = < 0.0001; Fig. 4f-h). This was associated with decreased axon cross-sectional area (*t*(383) = 2.185, *p* = 0.030; Fig. 4i) and reduced inner-to-outer diameter ratio of myelinated axons (‘G-ratio’; *t*(383) = 7.531, *p* = < 0.0001; Fig. 4j). This MC38-related decrease in G-ratio extended to myelinated axons of all calibers (F_7, 377_ = 11.05, *p* = < 0.0001; Fig. 4k-l). Sporadic abnormalities were also detected in unmyelinated axons (Supplementary Fig. 6b-c). Although the average number of unmyelinated axons per Remak bundle did not significantly differ (Supplementary Fig. 6d), unmyelinated axons in MC38 mice had a rightward shift in their distribution of cross-sectional area, indicative of axonal swelling (*t*(1694) = 13.28, *p* = <0.0001; Supplementary Fig. 6e-f).

**Figure 4.**
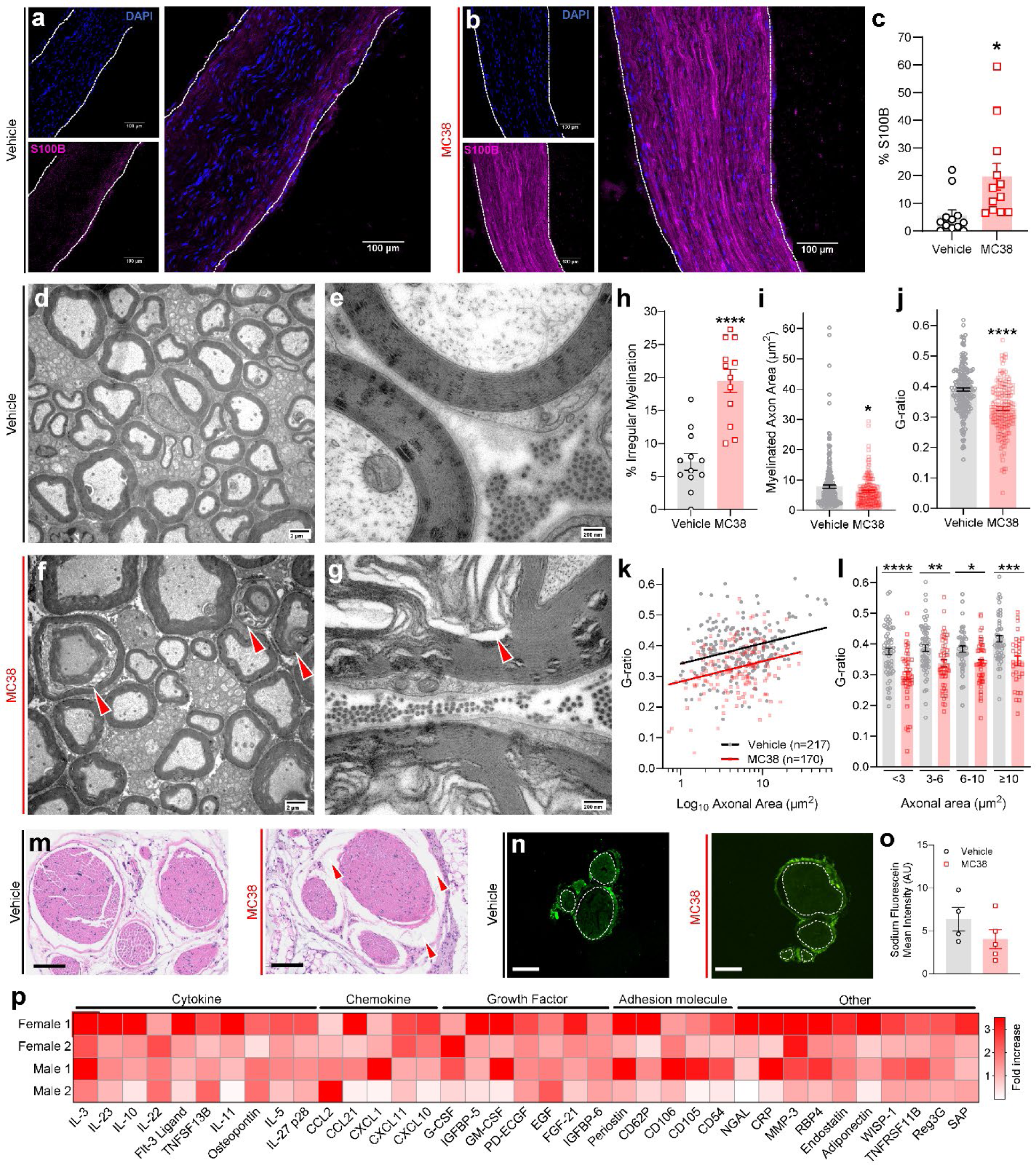
Schwann cell injury/demyelination and inflammation in peripheral nerves of MC38 tumor-bearing mice. (**a**) S100B staining density was low-to-undetectable in nerve samples from vehicle-treated mice, (**b-c**) but was significantly increased in MC38 nerves. (**d-e**) Transverse TEM images of sciatic nerve from vehicle-treated mice show grossly normal myelination of large-caliber axons, typified by compact, electron-dense myelin. MC38 mouse nerves show numerous irregularities in myelination (red arrowheads; **f-g**), with a significant increase in the percentage of myelinated axons surrounded by decompacted myelin (**h**). Combined with a small but significant decrease in average cross-sectional area of myelinated axons (**i**), this leads to a significant decrease in G-ratio (**j**). Error bars = SEM. */**** *p* = <0.05 / 0.0001, MC38 versus vehicle, unpaired T-test. This decrease in G-ratio is statistically significant across axons of all calibers (**k-l**), Error bars = SEM. */**/***/**** *p* = <0.05 / 0.01 / 0.001 / 0.0001, MC38 versus vehicle, one-way ANOVA, Sidak’s multiple comparisons test. (**m**) Transverse sections of sciatic nerve from vehicle-treated mice do not show any histological signs of neuroinflammation. Samples from MC38-injected mice show evidence of subperineurial edema (red arrowheads), without any obvious increase in cellular density or edema in the epineurium or endoneurium. Scale bar = 100 µm. (**n**) Representative images of sodium fluorescein extravasation into the endoneurium (dotted white lines), quantified in (**o**). Scale bar = 100 µm. (**p**) Semi-quantitative antibody array analysis of sciatic nerve homogenates from four MC38 mice (n=4, 2 males + 2 females) show widespread and variable upregulation of numerous pro-inflammatory cytokines, chemokines, growth factors and other inflammatory mediators when compared to vehicle-injected controls.

Consistent with the ultrastructural changes in the sciatic nerves of MC38 mice, histological stains of sciatic nerves revealed some evidence of epineurial edema, although individual fascicles and the surrounding perineurium remained intact (Fig. 4m). We did not detect increased sodium fluorescein extravasation into the endoneurium of MC38 mouse nerves, suggesting that tumor growth does not cause overt loss of blood-nerve barrier integrity (Fig. 4n-o). Messenger RNA for several cytokines and chemokines was elevated in sensory ganglia from MC38 mice^11^. Consistent with this, several cytokines, chemokines, growth factors and adhesion molecules were substantially upregulated (≥ 1.5-fold) in MC38 mouse nerves versus controls (Fig. 4p and Supplementary Table 3). This suggests MC38 tumor growth is associated with plasma extravasation in the epineurium and upregulation of inflammatory mediators that may facilitate leukocyte infiltration.

We next investigated the macrophage content of sciatic nerves and lumbar sensory ganglia in MC38 mice. We detected increased CD68 staining density in the epineurium (*t*(50) = 2.568, *p* = 0.013) and peri/endoneurium (*t*(50) = 3.148, *p* = 0.003) of sciatic nerves of MC38 mice (Fig. 5a-d). This was mirrored in the meninges and parenchyma of lumbar DRGs (*t*(30) = 2.395, *p* = 0.023 [meninges]; 4.387, *p* = 0.0001 [parenchyma]; Fig. 5e-h). Although we did not detect increased IgG deposition in MC38 sciatic nerves, NF200 expression was increased (*t*(21) = 2.876, *p* = 0.009; Fig. 5i-l). In contrast, NF200 expression was not increased in MC38 mouse DRGs, but we detected increased IgG immunoreactivity (*t*(20) = 2.596, *p* = 0.017; Fig. 5m-p). This suggests the inflammation and macrophage accumulation in the peripheral nervous system of MC38 mice may be associated with autoantibody deposition in sensory ganglia.

**Figure 5.**
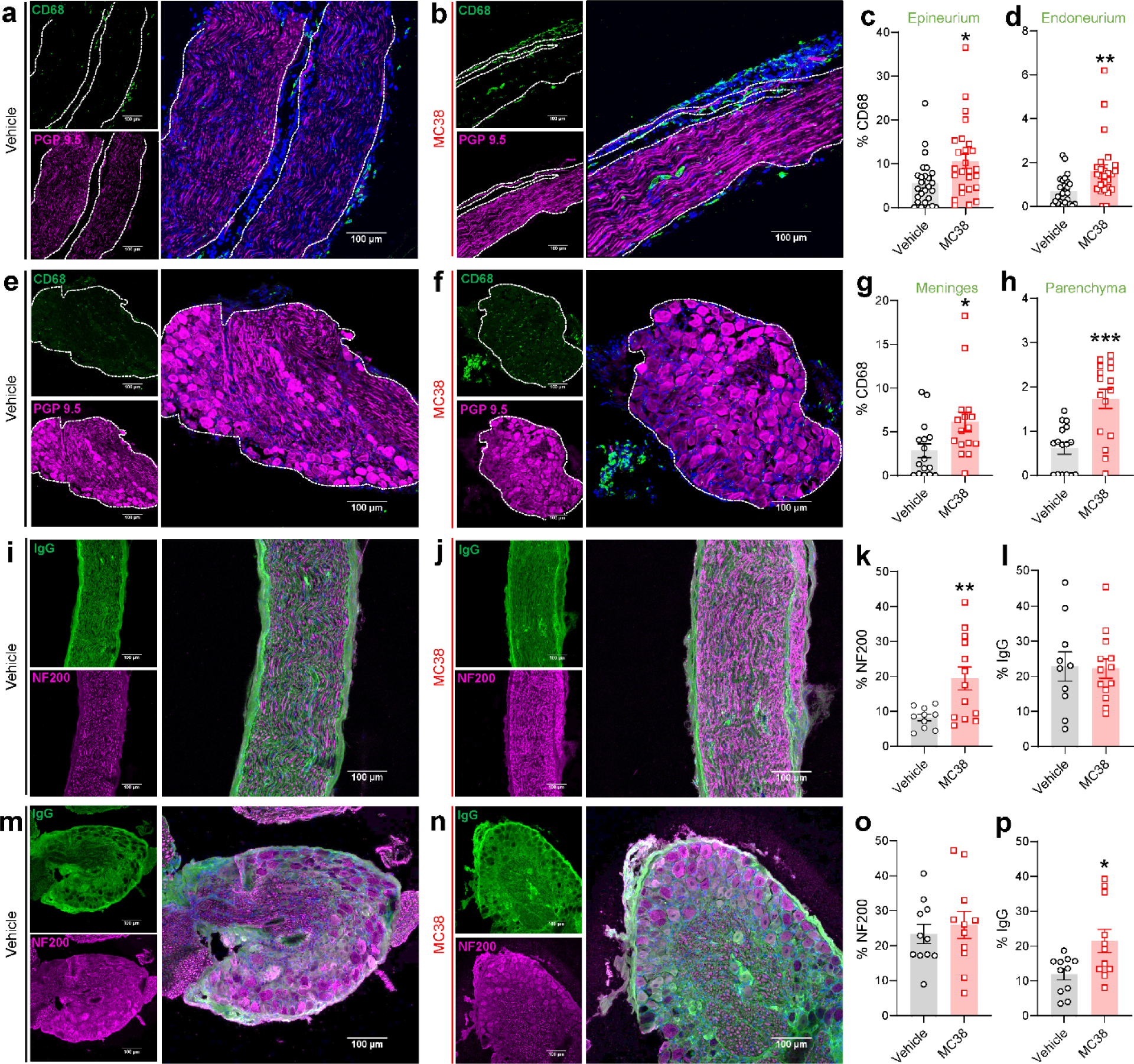
Macrophage infiltration and IgG deposition in the peripheral nervous system in MC38 tumor mice. (**a**) Representative images of longitudinal sciatic nerve sections from vehicle and (**b**) MC38 mice show a significant increase in endoneurial and epineurial CD68 staining density (green), quantified in (**c-d**). Dotted white lines denote the border between PGP9.5-positive axons and epineurium. DAPI = blue. (**e-h**) Similarly, dorsal root ganglia from MC38 mice show significantly increased CD68 density in the parenchyma (i.e., around neuronal somata), as well as in the meninges. Dotted white lines denote the border between PGP9.5-positive somata and meninges. (**i**) Representative image of sciatic nerve from vehicle and (**j**) MC38-injected mice, stained with antibodies against IgG and myelinated axon marker NF200. (**k**) A significant increase in sciatic nerve NF200 immunoreactivity was detected, (**l**) but not IgG. (**m**) Dorsal root ganglia from vehicle and (**n**) MC38 mice show no increase in NF200 intensity (**o**), but a significant increase in IgG staining density in MC38 tumor mice (**p**). Error bars = SEM. */**/*** *p* = <0.05 / 0.01 / 0.001, MC38 versus vehicle, unpaired T-test.

### Ryanodine receptor oxidation is associated with DRG neuron dysfunction in MC38 mice

In addition to dyslipidemia, inflammation and cytokine release induce calcium dysregulation via ryanodine receptors (RyR)^27^. We detected a significant upregulation of *Ryr1* mRNA in MC38 mouse DRG in our prior dataset^11^ (Supplementary Fig. 6a). Consistent with inflammation-associated oxidative stress, we detected increased RyR oxidation and reduced calstabin binding in MC38 mouse DRG, both indicators of dysfunctional ‘leaky’ RyR (*t*(5) = 2.686, *p* = 0.044; Fig. 6a-b). These deficits were associated with elevated serum TGF-β1 (*t*(8) = 2.522, *p* = 0.036; Fig. 6c), as has been described for skeletal muscle RyR dysfunction secondary to bone metastasis^27^.

**Figure 6.**
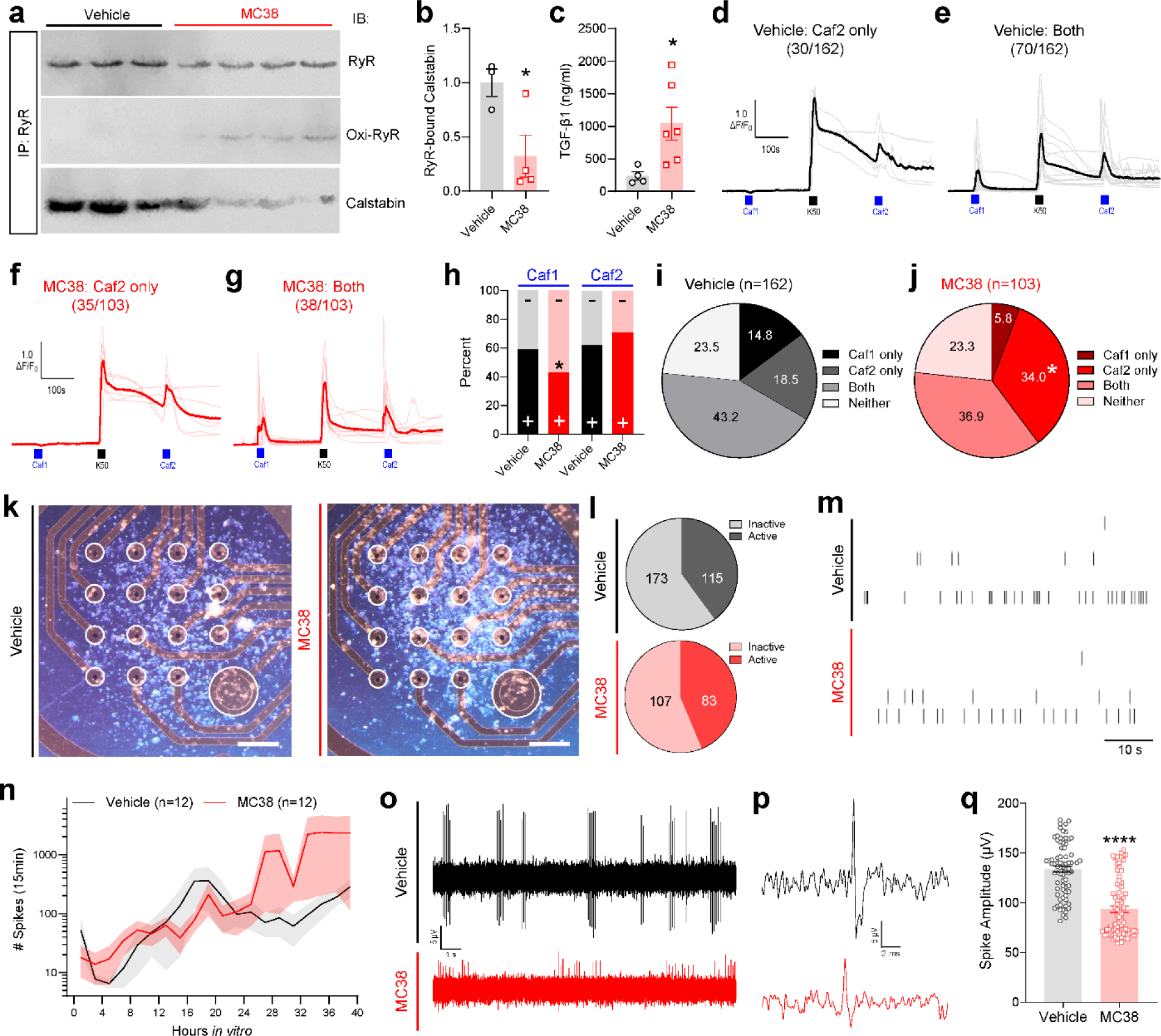
Ryanodine receptor oxidation, calcium dyshomeostasis and reduced spike amplitude in DRG neurons from MC38 tumor-bearing mice. (**a**) Ryanodine receptor (RyR) complexes were immunoprecipitated from lysates of dorsal root ganglia. An increase in oxidized RyR was associated with a reduction in RyR-bound calstabin (**b**) and elevated plasma TGF-β (**c**). *=p<0.05, MC38 versus vehicle, unpaired T-test. (**d-e**) Live-cell Ca^2+^ imaging shows that sensory neurons from vehicle-injected mice vary in their endoplasmic reticulum (ER) Ca^2+^ content. 20 mM caffeine mobilizes calcium from ER stores by opening RyR complexes. Thirty out of 162 (18.5%) vehicle-injected mouse neurons did not respond to the first pulse of 20 mM caffeine (‘Caf1’; blue bar, 15s), but did respond to the second caffeine pulse (‘Caf2’; blue bar, 15s) which was delivered three minutes after depolarization with 50 mM potassium chloride (‘K50’; black bar, 15s). A plurality of neurons (43.2 %) from vehicle-treated mice respond to both caffeine pulses, reflecting steady-state retention of Ca^2+^ within the ER. Gray traces represent individual neurons, thick black traces represent the mean response. (**f-g**) In contrast to vehicle-treated mouse neurons, more neurons from MC38-injected mice fail to respond to Caf1 (34.0 %), consistent with ER Ca^2+^ depletion. Pink traces represent individual neurons, thick red traces represent the mean response. (**h**) Significantly fewer neurons from MC38 mice respond to Caf1. The response to Caf2 was normalized by K50 depolarization, such that Caf2 response rates do not differ between vehicle and MC38 mice. ‘+’ = response, ‘- ‘ = no response. (**i, j**) A significant increase in the proportion of ‘Caf2 only’ neurons (i.e., did not respond to Caf1, but did respond to Caf2) in MC38-injected mice. * = *p* <0.05, two-sided Fisher’s exact test. (**k**) Dissociated DRG neurons are similarly viable, dense, and adherent in cultures prepared from vehicle-injected and MC38 tumor-bearing mice, 48 hours after plating onto multi-electrode array plates. Scale bar = 0.2 mm. (**l**) at 24h *ex vivo*, vehicle- and MC38-injected mouse DRGs (3 weeks post-injection) register similar proportions of active electrodes across all wells (n=16 electrodes per well). (**m**) Representative raster plots showing typical spontaneous activity (suprathreshold spikes) across 60 seconds and five electrodes in dissociated DRG neurons from vehicle control (upper) and MC38 (lower) mice. (**n**) Quantification of average spike number per well in cultures of dissociated vehicle and MC38 mice from 1 to 40 hours *in vitro*. No significant differences in activity level were noted. (**o**) Representative 10s and (**p**) 10ms voltage traces, recorded at 19h *ex vivo* show reduced spike amplitude in MC38 DRG neurons compared to vehicle controls, quantified in (**q**). Error bars = SEM. **** = *p* <0.0001, MC38 versus vehicle, unpaired T-test.

To determine whether RyR oxidation affected neuronal function, we carried out live-cell Ca^2+^ imaging on dissociated DRG neurons. RyR stimulation with caffeine elicited Ca^2+^ flux in 58% of neurons from vehicle control mice (94/162 neurons; Fig. 6d-e), reflecting Ca^2+^ mobilization from the endoplasmic reticulum (ER) into the cytosol. Initial caffeine responses were significantly less frequent in MC38 mouse neurons (43%; 44/103 neurons; *p* = 0.034, Fisher’s exact test, Fig. 6f-h), suggesting that ER Ca^2+^ stores are depleted in MC38 mouse neurons. Accordingly, KCl-stimulated influx of extracellular Ca^2+^ did not change caffeine response frequency in control neurons, but it did normalize the response deficit in MC38 neurons (Fig. 6h), suggesting Ca^2+^ uptake into the ER is unaffected in MC38 tumor mice. The frequency of neurons responding to caffeine only after KCl (34.0%) was significantly greater in MC38 mouse neurons than control (18.5%; χ^2^(3) = 11.25, *p* = 0.011; Fig. 6i-j). Within the subset of neurons that responded to the first pulse of caffeine, Ca^2+^ amplitudes did not significantly differ between vehicle controls and MC38 mice. A slight but significant reduction in amplitude of the second caffeine pulse in MC38 mice contributed to a reduced pulse 2: pulse 1 amplitude (*t*(172) = 2.166, *p* = 0.032; Supplementary Fig. 7a-c). These observations are broadly consistent with depletion of ER Ca^2+^ due to TGF-β-driven oxidization of neuronal RyR.

We next used multi-electrode arrays to determine whether dysregulated ER Ca^2+^ homeostasis affected sensory neuron activity in MC38 mice. At 48 hours *ex vivo*, DRG neurons from vehicle and MC38-injected mice showed similar levels of neuronal density (Fig. 6k). Twenty-four hours after dissociation, vehicle and MC38 mouse neurons showed a similar number of active electrodes (>1 spike/min; Fig. 6l) and spikes per electrode (Fig. 6m-n), suggesting that sensory neuron viability and spontaneous activity were not adversely affected by MC38 tumor growth. Similarly, the frequency of electrically-evoked spikes did not significantly differ (Supplementary Fig. 7d-j). However, spikes in MC38 mouse neurons were of significantly lower amplitude (*t*(157) = 9.049, *p* = < 0.0001; Fig. 6o-q). Collectively, this suggests RyR oxidation is associated with a deficit in spike amplitude rather than frequency, which could underlie any sensory/proprioceptive deficits associated with MC38 tumor growth.

### Inflammation and nerve degeneration in nerves of rhesus macaques with CRC

To test the generalizability of our findings in MC38 mice, we analyzed sciatic nerves of rhesus macaques with a diagnosis of CRC at necropsy, compared with those that did not receive a cancer diagnosis. Control nerves show few signs of inflammation or nerve damage (Fig. 7a-b, d-e) and only sporadic Iba1 immunoreactivity (a marker of macrophages and other cells of histiocytic origin; Fig. 7c, f). However, cellular infiltrates, axonal degeneration, and increased Iba1 immunoreactivity (indicating an increased histiocytic infiltrate) were readily detectable in CRC macaques (Fig. 7g-l). This was reflected in a significant difference in the frequency and degree of inflammatory cell infiltrate (Infiltrate score ≥2: 2/9 controls, 16/18 CRC, *p* = 0.002, Fisher’s exact test, Fig. 7m) as well as degeneration (score ≥3: 1/9 controls, 11/18 CRC, *p* = 0.031, Fisher’s exact test, Fig. 7n). Accordingly, Büngner bands (regenerating nerve tracts formed by Schwann cells following axonal degeneration) were detected in 1/7 control samples, but 11/15 CRC samples (*p* = 0.02, Fisher’s exact test, Fig. 7o). These data indicate that several key neurological abnormalities detected in MC38 tumor-bearing mice are recapitulated in spontaneous non-human primate cases of CRC.

**Figure 7.**
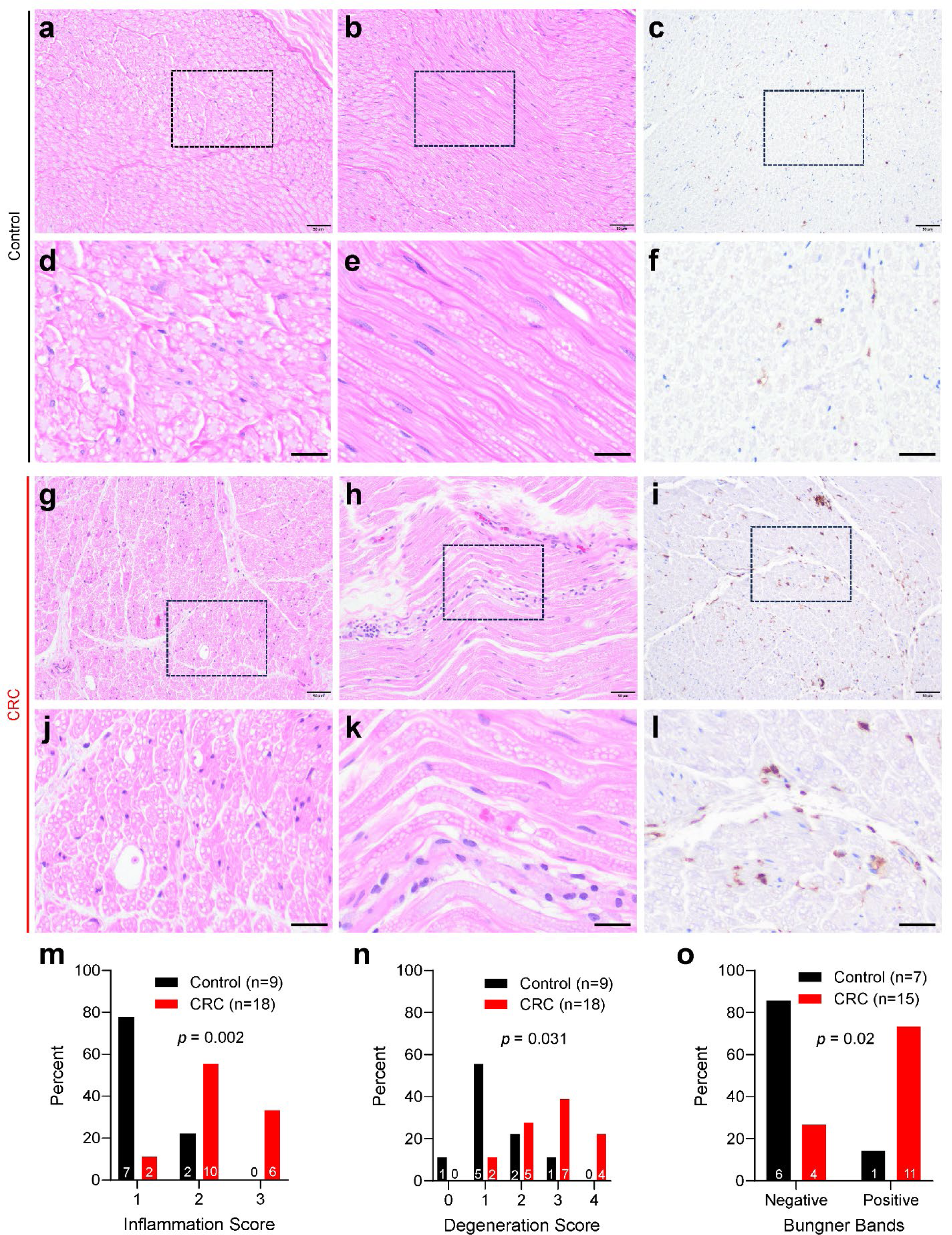
Peripheral nerve inflammation in mismatch repair-deficient CRC in rhesus macaques. (**a**) transverse and (**b**) longitudinal H&E micrographs of sciatic nerve from non-cancer control rhesus macaques. Dotted lines show insets in (d-e). (**c, f**) Iba1 immunohistochemistry shows sparse labeling typical of healthy resident macrophage density. Scale bars = 50 µm (**a-c**), 25µm (**d-f**). (**g**) transverse and (**h**) longitudinal sections of sciatic nerve from CRC macaques show evidence of cellular infiltration and axonal degeneration with digestion chambers (**g, j**) and regeneration (Büngner bands (**h, k**)), along with a notable increase in Iba1 immunoreactive histiocytic cells (**i, l**). Semi-quantitative analysis demonstrates a significant increase in inflammation score (**m**), degeneration score (**n**) and the presence of Büngner bands (**o**). *p* = <0.05, Fisher’s exact test.

## Discussion

Increasing cancer survivorship inevitably places greater emphasis on long-term complications. The widespread and debilitating neurologic complications in cancer survivors lack effective therapies, necessitating an improved understanding of somatosensory nervous system damage due to cancer. For example, paraneoplastic neurologic syndromes are regarded as an infrequent consequence of cancer-driven autoimmunity. Diagnoses of ‘true’ paraneoplastic neuropathy in CRC are rare^28^, but prior studies have uncovered deficits in IENF density and sensory acuity in small cohorts of CRC patients^9, 10^. Because these were unanticipated findings, these studies were not designed to address causality. For example, type 2 diabetes and excessive alcohol intake are potential confounds which cause neuropathy and elevate the risk of developing CRC^29, 30^. Therefore, our objective was to characterize CRC-driven neuropathy in preclinical models to address causality, and to identify potential causative factors. Our results using an orthotopic, immunocompetent mouse model and spontaneous rhesus macaque model of CRC suggest there is indeed a causal link between CRC and development of sub-clinical neuropathy.

Consistent with our prior observations in male mice^11^, IENF density was reduced in females, in the absence of any overt hypersensitivity. This sensory neuropathy without evidence of tactile hypersensitivity parallels non-painful neuropathy in CRC patients, where touch detection thresholds were impaired^10^. Interestingly, this study also identified warming and cooling detection deficits, which we did not detect in mice in the thermal gradient ring test. The increased levels of IL-10 in sciatic nerves from MC38 mice may explain why this neuropathy is not associated with pain; prior studies associated elevated levels of anti-inflammatory cytokines such as IL-10 with non-painful neuropathy, and downregulation in patients with painful neuropathy^31^. In fact, the most common first manifestation of paraneoplastic and other neuropathies is sensory loss^32^, which may contribute to underdiagnosis. Similarly, the loss of IENFs is often detected in diabetic neuropathy patients with non-painful neuropathy^33^. The lack of robust, sustained hypersensitivity in MC38 tumor-bearing mice following naltrexone suggests that MC38 mice do not experience substantial latent pain sensitization. This contrasts with cisplatin-induced neuropathy, which shows rapid and robust naltrexone-induced reinstatement of pain in the remission phase^12^.

Because the mechanisms underlying different forms of neuropathy vary widely, the persistence of CRC neuropathy and the degree to which it transitions to painful neuropathy remain to be established. However, we detected motor deficits in gait analysis and balance beam tasks. Our prior study in this model did not detect significant sensory loss in MC38 mice^11^, perhaps suggesting these deficits are driven by proprioceptor/motor neuron deficits rather than overt sensory loss. A prior report using a motor coordination task in CRC rats saw similar deficits following chemotherapy^34^, which could provide a basis for the increased fall risk seen in cancer survivors, independent of age or medication use^35^.

In search of factors underlying CRC neuropathy, we present evidence that metabolic and hemostasis markers in MC38 mice remained largely normal, with the notable exception of lipids and fibrinogen. An increased clotting propensity is consistent with CRC patient data^36^ and our prior analysis of DRG from MC38 tumor mice^11^. While the underlying mechanism (if any) that links hypercoagulability to CRC-induced neuropathy remains to be determined, it is conceivable that coagulation impairs endoneurial blood flow, leading to ischemic neuropathy^17^.

Neurodegenerative diseases such as multiple sclerosis are accompanied by plasma lipid dysregulation^22^, and the same is true for plasma, tumor and adipose lipids in CRC and other malignancies^21, 37^. Most cancers promote FA synthesis, elongation and desaturation to support their growth^38^. Accordingly, PC/LPC, PE, and PI were elevated MC38 mouse plasma. We also detected enrichment of cholesterol esters, which has been reported in lung cancer^37^, and depletion of short chain FAs and enrichment of long chain FAs, as well as an increase in PUFAs, alongside depletion of monounsaturated/unsaturated FAs^39^. Sciatic nerve lipids mirrored these systemic changes; we detected hexosylceramide, sphingomyelin and LPC enrichment and triglyceride depletion. Importantly, these disruptions parallel the dysregulation seen in nerve injury and neurodegeneration^40, 41^. It will be instructive to establish the extent to which peripheral nerve lipid dysregulation underlies both inflammation and neuropathy. Considering the RyR dysfunction in the ER of MC38 mouse neurons, it is tempting to speculate that cancer-induced ER stress either initiates or sustains dyslipidemia, since activation of the unfolded protein response leads to many similar derangements in lipid metabolism^21^. Additionally, numerous links between CRC, gut dysbiosis and lipid metabolite profile^42^ could hint at a mechanism for CRC-induced neuropathy that is driven by changes to gut microbiota.

Macrophage accumulation and increased NF200 expression are hallmarks of inflammatory neuropathy in other disease models^14, 43^. Because macrophages adopt protective as well as pathological roles in the peripheral nervous system^44^, establishing the functional impact of macrophage accumulation will be important. In addition to leukocyte infiltration and dysregulation of lipid metabolites, Schwann cell secretion of S100B and myelin decompaction are also symptoms in other neuropathies e.g. following cisplatin chemotherapy^45^. The dysregulation of myelin across all calibers of myelinated axons suggests motor axons as well as sensory Aβ and Aδ fibers responsible for transducing innocuous touch and proprioceptive stimuli are all impacted. Furthermore, prior studies suggest that swelling of unmyelinated axons occurs in response to injury in neuropathy^46^. It is possible all these signs of dysfunction are secondary to pro-inflammatory cytokine production or circulating dyslipidemia^47^. In either case, dysfunctional myelination is likely connected to the observed motor coordination deficits, because compromised myelination reduces action potential velocity and amplitude in other neuropathies^48^.

Traditionally, paraneoplastic neurologic syndromes are represented by autoimmunity against the nervous system, caused by tumor-driven production of ‘onconeural’ antibodies. Consequently, signs and symptoms of paraneoplastic neurologic syndromes resemble other autoimmune neurologic conditions, such as complex regional pain syndrome, multiple sclerosis and Guillain-Barré syndrome^49^. Paraneoplastic neuropathy is most commonly associated with lung cancer and lymphoma, but we detected IgG deposition in the DRG of MC38 tumor-bearing mice that is consistent with antibody production against neuronal antigens seen in other, more prototypical paraneoplastic neuropathies. Therefore, IgG deposition may underlie at least some of the CRC-related inflammation and neuropathy, though further work is needed to understand the antigen(s) involved and the relative contribution of autoantibodies to CRC-driven neuropathy.

It is also plausible that the degree of inflammation is influenced by the mutational burden and therefore immunogenicity of different CRC cases. Notably, both MC38 cells and the rhesus macaques used in this study are models of mismatch repair-deficient/microsatellite instability CRC. This is due to mutations in *Msh3* and *Pold1* in MC38 cells, and *MLH1* germline mutation or hypermethylation in rhesus macaques^50, 51^. This leaves open the question of whether mismatch repair-proficient/microsatellite stable CRC causes neuropathy via similar mechanisms, though our observations using CT26 tumors in mice suggest neuropathy also develops in that context^11^. Nevertheless, thorough exploration of the relationship between mutational burden, neoantigen expression and neuropathy is clearly warranted. Interestingly, inflammatory bowel disease has previously been linked to neuropathy^52^. Since chronic inflammatory conditions such as ulcerative colitis substantially increase CRC risk^53^, it is conceivable that many CRC patients could be developing neuropathy years prior to a CRC diagnosis.

Malik et al.^28^ report that symptomatic neuropathy is rare in CRC, but our findings add to the weight of evidence that the true incidence of neuropathy in CRC may be under-reported. The authors of this study also speculated that the rare instances where CRC was tied to a clinical diagnosis of paraneoplastic neuropathy most closely resemble chronic inflammatory demyelinating polyneuropathy (CIDP), given the macrophage infiltration of nerves, phagocytosis of myelin and autoantibody deposition^54^. Corticosteroid therapy is often effective in CIDP, although the risk versus benefit of immunosuppression in patients being treated for CRC remains to be established. Ultimately, these data raise the prospect that CRC-driven neuropathy represents an underappreciated risk factor for development of neurologic issues in CRC survivors. It is possible that subacute CRC neuropathy could interact with chemotherapy-induced neurotoxicity or immune-related toxicities associated with immune checkpoint inhibitors, though our understanding of the links between immunotherapies and chronic neuropathic pain is still in its infancy. Nevertheless, pre-existing neuropathy is a major risk factor for development of chemotherapy-induced peripheral neuropathy^6^. The latent neuropathy associated with CRC could therefore represent a ‘priming’ event in susceptible individuals, augmenting the pain response to additional insults. Future studies should address any potential relationship between CRC-induced neuropathy and the development and resolution of pain in response to subsequent insults.

## Materials and Methods

### Animals

All experiments and procedures involving the use of live animals were approved by MD Anderson Cancer Center Animal Care and Use Committee (Houston, TX, USA), in accordance with the National Institutes of Health’s Guide for the Care and Use of Laboratory Animals. C57BL/6J mice (Strain #: 000664) were purchased from Jackson Laboratories, randomly assigned to experimental groups and housed in the MD Anderson Cancer Center’s Research Animal Support Facility on a 12-h light cycle with access to food and water *ad libitum*. Mice were used between 8-12 weeks of age.

A closed breeding colony of Indian-origin rhesus macaques (*Macaca mulatta*) bred at the MD Anderson Cancer Center Michale E. Keeling Center for Comparative Medicine and Research since 1989 spontaneously develops CRC with microsatellite instability and mismatch repair deficiency. The colony is specific pathogen free for Herpes B, Simian retroviruses and *Mycobacterium tuberculosis.* A subset of these animals (approximately 5%^51^) harbor a pathogenic germline mutation in *MLH1* (c.1029C<G, p.Tyr343Ter)^50, 51^. Animals in the colony that die or are euthanized for meeting clinical endpoints are submitted for complete diagnostic necropsy including gross examination and histopathology. Review of pathology records identified animals diagnosed with CRC at necropsy and controls with no CRC from which archived FFPE sciatic nerve was available. Eighteen cases of CRC (n=14 females, 4 males, average age 21.3 ± 2.9 years) were compared against nine non-CRC controls (n=8 females, 1 male, average age 16.7 ± 5.8 years). This sex distribution is representative of the colony (∼80% female). Exclusion criteria were age <5 years (pre-sexual maturity), diagnosis of other systemic inflammatory conditions (e.g. sepsis, rheumatoid arthritis), seropositivity for *Trypanosoma cruzi*, and autolysis of tissues at necropsy.

### Orthotopic MC38 Model and Bioluminescent imaging

The MC38 colon adenocarcinoma cell line was obtained from Alstem Cell Advancements (MC38-Luc-RFP; RRID: CVCL_B7NQ). Cells were maintained in 10 cm TC dishes at 37 °C, 5% CO_2_ in high glucose DMEM with 10% fetal bovine serum, 2 mM glutamine, 0.1 mM nonessential amino acids, 10 mM HEPES, and 1x penicillin/streptomycin. Cells were used within 10-18 passages. Orthotopic tumors were implanted as previously described^11^. Briefly, 2×10^5^ MC38 cells in 50 µl growth medium were transanally injected into the rectal submucosa under isoflurane anesthesia.

Approximately 95% of injections resulted in successful engraftment. Mice without detectable bioluminescent signal two weeks post-injection were excluded from analysis. Unless otherwise stated, experiments were carried out on mice three weeks after vehicle/MC38 cell injection, since this was when significant peripheral neuropathy first became detectable^11^.Tumor growth was monitored weekly for up to 4 weeks by bioluminescent imaging using an IVIS Lumina XR (PerkinElmer), as described previously^11^. Briefly, mice were injected with D-luciferin (Goldbio; 150 mg/kg, i.p. in PBS) and imaged 10 minutes post-injection using auto-exposure settings.

Luminescence was quantified within a circular ROI overlying the colorectal region and expressed as average radiance (p/s/cm^2^/sr) using Living Image software (PerkinElmer).

### Behavioral Assays

Mice were habituated to all behavior testing equipment 24 hours prior to beginning testing. Testing was split during the light cycle, and carried out during the same time of day, either between 07:00h and 11:00h or between 13:00h and 16:00h.

#### Von Frey

Hindpaw mechanical sensitivity was determined using von Frey filaments in the ‘up-down’ method, as described previously^11^. Fifty percent paw withdrawal thresholds were calculated from the series of withdrawal responses. Naltrexone hydrochloride (Millipore Sigma) was dissolved in sterile PBS and injected subcutaneously into the nape of the neck at 3 mg/kg.

#### Burrowing

Burrowing activity is a pain-depressed behavior in mice that can be reversed by known analgesics, as we previously described^13^. Briefly, 30g of new bedding (¼’’ corn cob bedding) and 5 g of home cage bedding was placed in a 50 mm (diameter) x 125 mm (length), close-ended tube made from translucent red acrylic. Mice were placed in a cage individually and the bedding displaced from this tube after 15 minutes was calculated.

#### Thermal Gradient Ring

To detect any potential changes in temperature preference/sensitivity, mice were tested using a Thermal Gradient Ring device (Ugo Basile) attached to a PC running ANY-maze tracking software (Stoelting). Each mouse was allowed to freely explore the circular arena of the device, the floor of which was set to a temperature gradient, ranging from 15°C to 40°C. Mice were placed into the device at an innocuous temperature zone (approximately 23-27°C). After 15 minutes exploration, A 15-minute recording period began, where the dwell time in each pair of twelve temperature zones (each with a plane angle of 15 degrees / 0.26 rad) was recorded using the device’s top-down camera. For quantification, these 12 temperature zones were binned into four quartiles: Q1 (zones ranging from 15-19°C), Q2 (19-25°C), Q3 (25-32°C) and Q4 (32-40°C).

#### Catwalk Gait Analysis

Footfall patterns and gait were analyzed using the Catwalk XT device and software (Noldus). Gait indices were averaged across both hindpaws for four compliant runs (duration between 0.5s and 5s, maximum variation in run speed ≤60%) for each mouse at each timepoint and are depicted as percent change relative to baseline.

#### Beam Walk

Mice were trained on three consecutive days to cross a 12 mm-wide rectangular beam (‘wide flat’), followed by a 6 mm-wide rectangular beam (‘narrow flat’) and a 6 mm-diameter cylindrical beam (‘round’). All three beams are 85 cm in length and placed at an approximate 15-degree incline. The time taken to cross, and the number of errors (paw slips, inversions) were determined by investigators blinded to treatment. Mice that fell from the beam or failed to complete the task within 60 seconds were assigned the cutoff time of 60 seconds.

### Histology

Mice were perfused transcardially with 1x PBS followed by 10% neutral buffered formalin. Sciatic nerve and colon were processed in an automated processor with an overnight routine protocol (dehydration to 100% ethanol, followed by xylene and paraffin) to produce formalin fixed paraffin embedded blocks (FFPE). After sectioning FFPE blocks at 5µm, sections were stained with hematoxylin and eosin in a HistoCore SPECTRA ST (Leica Biosystems). Slides were deparaffinized through xylene and ethanol before incubating in hematoxylin for 22min, followed by washes and eosin for 1 min 45s, dehydration through an ethanol series and xylene before mounting. Stained slides were digitized in an Aperio AT2 slide scanner (Leica Biosystems) at 20 × magnification. FFPE sections of rhesus sciatic nerves were stained with hematoxylin and eosin. Images were semi-quantitatively scored for inflammatory cell infiltrates and axonal degeneration by a board-certified veterinary pathologist who was blinded to group identity.

### Immunohistochemistry and Fluorescence Microscopy

Dorsal root ganglia, sciatic nerves and 3mm hindpaw plantar punch biopsies were collected following transcardial perfusion with 1x PBS and 4% paraformaldehyde (PFA) in phosphate buffer (PB), pH 7.4, as described previously^11^. At least 3 biological replicates/groups were measured from at least two different cohorts of mice for all tissues. Plantar tissues were post-fixed in 4% PFA overnight before being transferred into 15% sucrose and 10% EDTA in PBS for 48h for decalcification of bones, as described previously. Sciatic nerves and DRG were incubated in 15% sucrose in PBS without EDTA. After further overnight incubation in 30% sucrose, 40µm lateral plantar, 20µm sciatic, and 10µm DRG sections were collected onto microscope slides using a Leica CM3050S cryostat. Immunohistochemistry was also performed as previously described^11^. After three 5-minute washes with 0.1M PB, slides were blocked and permeabilized with blocking buffer (10% goat serum, and 0.3% Triton X-100 in 0.1M PB) for 1h at 4°C. Slides were then incubated overnight at 4°C in primary antibodies diluted in blocking buffer (rabbit anti-PGP9.5: Abcam, catalog #ab108986; 1:250; rat anti-mouse CD68 (clone FA-11): Fisher Scientific, catalog # NC9471873; 1:500, rabbit anti-S100B (clone CL2720): Abcam, catalog # AB227914; 1:100). The following day, samples were washed 3 times in blocking buffer and then incubated with secondary antibody (1:500 IgG-Alexa Fluor 488 / goat anti-rabbit IgG-Alexa Fluor 647 and goat anti-rabbit IgG-Alexa Fluor 594 antibodies in blocking buffer and 1µg/ml DAPI for 4h, protected from light at 4°C.

IgG staining followed the same procedure, but used donkey serum in the blocking buffer in place of goat serum, blocked tissues overnight with blocking buffer and stained with a polyclonal goat anti-mouse F(ab’)_2_ fragment (1:100; Jackson ImmunoResearch 115-005-072) and rabbit anti-mouse NF200 (1:100; Sigma N4142), which were detected using 1:500 donkey anti-goat IgG-Alexa Fluor 488 and 1:500 donkey anti-rabbit IgG-Alexa Fluor 555, respectively. Following secondary incubation, slides were washed 3 times in 0.1M PB, mounted with Fluorsave, coverslipped, sealed with nail polish and stored at −20°C. Images were taken using a Nikon Ti2 confocal microscope at 20x magnification. Images are a composite of 10 focal planes in a 40 µm z-stack at 2 µm increments. 3-4 different sections were imaged from the same animal. Following image acquisition, density of staining including IENFs was determined as described previously^11^ using Nikon NIS-Elements Imaging Software. Signal density was calculated by the application of ROIs guided by DAPI staining to identify borders between cutaneous/neuronal/meningeal structures.

Immunohistochemistry of rhesus sciatic nerves for Iba1 was conducted on 4µm sections of FFPE tissue adhered to positively charged slides. Slides were immersed in 95°C Target Retrieval Solution (Dako) for 20min, followed by a gradual cooling step. Slides were then immersed and rinsed in wash buffer. The next steps were followed sequentially by a rinse with wash buffer and performed on the Fisher Autostainer: block slides for endogenous peroxidase activity using dual enzyme block (Dako), Background Sniper (Biocare), Fc Receptor Block (Innovex), primary antibody (rabbit monoclonal anti-Iba1 (clone EPR6136(2)):Abcam, #ab178680; 1:300) for 30 minutes, Secondary polymer (Envison+, DAKO), 3,3’-diaminobenzidine substrate for 10 minutes (DAKO) and counter stained for 5 minutes with automated hematoxylin (DAKO). Slides were then dehydrated through alcohol gradients and xylene before being coverslipped.

Blood-nerve barrier permeability was assessed following intravenous injection of sodium fluorescein (20mg/kg). One hour after injection, sciatic nerves were collected and 5µm transverse sections were imaged on the Olympus System Microscope Model BX53.

### Transmission Electron Microscopy

Samples were fixed using 3% glutaraldehyde and 2% paraformaldehyde in 0.1M cacodylate buffer (pH 7.3). Samples were then washed in 0.1M sodium cacodylate buffer and treated with 0.1% Millipore-filtered cacodylate buffered tannic acid, post-fixed with 1% buffered osmium and stained *en bloc* with 0.1% Millipore-filtered uranyl acetate. Samples were dehydrated through an ethanol series and infiltrated and embedded in LX-112 medium. Samples were then polymerized in a 60°C oven for approximately three days. Ultrathin sections were cut using a Leica Ultracut microtome and stained with uranyl acetate and lead citrate in a Leica EM Stainer. Stained samples were examined in a JEM 1010 transmission electron microscope (JEOL USA) using an accelerating voltage of 80 kV. Digital images were obtained using an AMT imaging system (Advanced Microscopy Techniques). Morphometric analysis of cross-sectional diameter/area was conducted using NIH ImageJ 1.54h.

### Lipidomics

Plasma and sciatic nerve samples for lipidomic analysis were snap-frozen on dry ice. Tissues from two mice were pooled together for each technical replicate. Blood was collected via cardiac puncture, collected in Precellys tubes and centrifuged at 1500 x g for 15 minutes at 4°C. 20µL of plasma was used for untargeted lipidomics analysis. Plasma samples were extracted by adding 200µL of extraction buffer (ice-cold ethanol containing 1% 10mM butylated hydroxytoluene, and 2% Avanti SPLASH^®^ LIPIDOMIX^®^ Mass Spec Standard). Samples were vortexed for 10min, placed on ice for 10min, and centrifuged at 17,000g for 10 min at 4°C. Supernatants were then collected for LC-MS analysis. Nerves were pulverized and then homogenized in ethanol, vortexed for 5min, placed on ice for 10min, and centrifuged at 17,000g for 10min at 4°C. The supernatants were evaporated to dryness under nitrogen and 200 µL of extraction buffer was added to reconstitute. Mobile phase A (MPA) was 40:60 acetonitrile:water with 0.1% formic acid and 10mM ammonium formate. Mobile phase B (MPB) was 90:9:1 isopropanol:acetonitrile:water with 0.1% formic acid and 10mM ammonium formate. The chromatographic method included a Thermo Fisher Scientific Accucore C30 column (2.6µm, 150 x 2.1mm) maintained at 40°C, a mobile phase flow rate of 0.200mL/min, an autosampler tray chilling at 8°C, and a gradient elution program as follows: 0-3min, 30% MPB; 3-13min, 30-43% MPB; 13.1-33min, 50-70% MPB; 33-48min, 70-99% MPB; 48-55min, 99% MPB; 55.1-60min, 30% MPB. The injection volume was 10 µL. A Thermo Fisher Scientific Orbitrap Fusion Lumos Tribrid mass spectrometer with heated electrospray ionization source was operated in data dependent acquisition mode, in both positive and negative ionization modes, with scan ranges of 150 –1500m/z. An Orbitrap resolution of 240,000 (FWHM) was used for MS1 acquisition and spray voltages of 3,600 and −2900V were used for positive and negative ionization modes, respectively. Vaporizer and ion transfer tube temperatures were set at 275°C and 300°C, respectively. The sheath, auxiliary and sweep gas pressures were 35, 10, and 0 (arbitrary units), respectively. For MS2 and MS3 fragmentation a hybridized HCD/CID approach was used. Data were analyzed using Thermo Scientific LipidSearch software (version 5.1) and R scripts written in-house. The peak area (raw relative abundance) of each metabolite was normalized by dividing by the tissue weight of the respective sample followed by z-score normalization.

### DRG neuron culture

DRG neurons were isolated from adult mice and enzymatically digested with collagenase and pronase before plating onto poly-L-ornithine and laminin-coated glass coverslips or multi-electrode array plates. To remove excessive debris, cell suspensions were centrifuged through a 12.5% and 28% Percoll density gradient before resuspension in culture media, as previously described^11^. Cells were cultured in a 1:1 ratio of TNB media:protein-lipid complex, and DMEM with 10% fetal bovine serum at 37°C in a 5% CO_2_ incubator for 24h before use in calcium imaging experiments, and for 1h before use in multi-electrode array recordings.

### Calcium Imaging

Calcium imaging of mouse DRG neurons was carried out as previously described^11^, with minor modifications. Dissociated neurons were plated onto poly-L-ornithine and laminin-coated 12mm circular glass coverslips. At 24h *in vitro*, individual coverslips were incubated with 2µM Fura-2 AM (Invitrogen) at room temperature for 20min. Fura-2 dye was dissolved in HEPES-buffered HBSS (‘HH buffer’), containing (in mM): 140 NaCl, 5 KCl, 1.3 CaCl_2_, 0.4 MgSO_4_, 0.5 MgCl_2_, 0.4 KH_2_PO_4_, 0.6 NaHPO_4_, 3 NaHCO_3_, 10 glucose, 10 HEPES, adjusted to pH 7.4 with NaOH and 310 mOsm with sucrose. Coverslips were then mounted into an MS-512SP recording chamber (ALA Scientific) on the stage of an inverted Nikon Ti2 microscope. Coverslips were continuously superfused with HH buffer at a flow rate of 1-2mL/min. Fura-2 dye was alternately excited at 340 and 380nm (2nm band-pass filter, 100ms exposure) at 1Hz using a Lambda LS Xenon light source and controller (Sutter Instruments). Fluorescence emitted at 510 nm was captured using a 10x/NA 0.5 objective and sCMOS pco.edge camera. Nikon NIS Elements software was used to calculate the F_340_:F_380_ ratio for each neuron. Because caffeine activates the sensory ion channel TRPA1, A967079 (selective TRPA1 antagonist, 1µM) was added to HH buffer throughout. After a 60s baseline reading, 20mM caffeine was pulsed for 15s, followed by 180s of recovery. This was followed by a 15s pulse of 50mM potassium chloride (‘K50’) to induce extracellular calcium influx. A second pulse of 20mM caffeine was applied 180s after potassium chloride, followed by a final 180s of HH buffer. The threshold for a calcium response during agonist application was defined by a peak F_340_:F_380_ ratio >5.5 standard deviations over baseline noise.

### Multi-Electrode Array

Dissociated DRG neurons were plated in 10µl droplets (∼10^4^ neurons) onto six poly-L-ornithine/laminin-coated wells of a 24-well CytoView MEA plate, as described previously^11^. After 1h to allow cell attachment, each well was filled with 0.75mL of culture media and the plate placed into a Maestro Edge MEA device (Axion BioSystems) at 37°C and 5% CO_2_. Real-time field potential recordings were acquired for a period of 15 minutes every two hours (12.5kHz sampling rate, 200-3000Hz band-pass per electrode across 16 electrodes per well). Spike number and firing rate were calculated using AxIS Navigator software and Neural Metrics Tool (Axion BioSystems). Evoked activity was delivered at 24h *in vitro* via four of the sixteen electrodes in each well using 3 stimulations at 10-second intervals. Biphasic stimulation was delivered at 0.3 and −0.3V (each 0.75ms; 1.5ms total), followed by stimulation artifact elimination (2ms discharge at 1µA, soft switch at 1ms; 3ms total). In all recordings, spikes were defined as voltages >5.5x the standard deviation of the noise for each individual electrode. Coincident artefacts (0.24ms window) and spikes <2ms post-stimulation were excluded from analysis using Axion Neural Metrics Tool. Spike waveforms were exported from raw traces using Axion Data Export Tool.

### Immunoprecipitation and Western Blotting

RyR was immunoprecipitated from DRG lysates with 2µg anti-RyR (Santa Cruz Biotechnology, sc-376507) in 0.5mL of modified RIPA buffer [20 mM Tris-HCl (pH 7.5), 250mM NaCl,1mM EDTA, 1% NP-40, 1mM Na_3_VO_4_, and Protease Inhibitor Cocktail (Cell Signaling #5871S)] for 4h at 4°C. Immune complexes were incubated with Protein G Sepharose^®^ 4 Fast Flow (GE Healthcare, #17-0618-01) overnight at 4°C, and the beads were washed three times with RIPA buffer. The immunoprecipitates were size-fractionated on 4-20% SDS-PAGE gels (for RyR) and 15% (for Calstabin) and transferred onto PVDF membranes for 2.5h at 200mA. Immunoblots were developed using anti-RyR and anti-Calstabin (Santa Cruz Biotechnology, sc-133067). To determine RyR oxidation, the carbonyl groups in the protein side chains were derivatized to 2,4-dinitrophenylhydrazone (DNP-hydrazone) by reaction with 2,4-dinitrophenylhydrazine. DNP signal was determined using a specific anti-DNP antibody (Millipore, MAB2223). All immunoblots were developed using the SuperSignal West Pico PLUS Chemiluminescent Substrate (Thermo Fisher 34577) and detected using an Odyssey system (LI-COR). The bait protein RyR acts as an internal control. Relative band intensity was quantified using NIH ImageJ.

### Cytokine Quantification

Cytokine content of sciatic nerve lysates was determined semi-quantitatively using a Proteome Profiler Mouse XL Cytokine Array (R&D Systems ARY028) according to the manufacturer’s instructions and as described previously^11^. For plasma TGFβ1, blood was collected from mice via cardiac puncture. Platelet-free plasma was isolated and TGFβ1 concentrations were determined using a commercially available Quantikine mouse TGF-β1 ELISA (R&D Systems DY-1679) according to the manufacturer’s instructions.

### Statistics

Von Frey and Catwalk data were compared between groups and across time using two-way, repeated measures ANOVA with Tukey’s multiple comparisons test. Beam walk data were analyzed using paired T-tests, all other behavioral tests used unpaired T-tests. Mixed-effects analyses were used in place of two-way ANOVA when datapoints were missing. Immunohistochemistry- and Western blot densitometry, ELISAs, hematological assays, spike amplitudes and endpoint fluorescence were compared using unpaired Student’s T-test. Fisher’s exact test was used to analyze calcium imaging and rhesus macaque histology data. Gaussian best-fit distributions and all statistical analyses were performed using Prism 10.1.2 (GraphPad).

## Acknowledgments

CMG, AMC, LS, IM, PLL, TAG, CLH and AJS contributed to experimental design, data acquisition, interpretation of results and manuscript development. BW, KV, EAK and FRC contributed to data acquisition. AJS contributed to funding acquisition. The authors declare that they have no competing interests. This work was funded by a Rita Allen Foundation Award in Pain (to AJS). The Small Animal Imaging, Histopathology, Metabolomics, and Electron Microscopy Core Facilities used in this study are core MD Anderson research resources supported by the NIH/NCI under award number P30CA016672. We wish to thank Kenneth Dunner and Nathan Fiore for technical assistance with transmission electron microscopy and IgG immunohistochemistry, respectively.

## Data Availability

The lipidomic and proteome profiler datasets that support the findings of this study are included in this published article (and its supplementary information files). All other data generated or analyzed during this study are available from the corresponding author upon reasonable request.

