## Supplementary Figures for "Inflammatory Neuropathy in Mouse and Primate Models of Colorectal Cancer"

### Supplementary Materials and Methods

#### *Blood Glucose*

Blood glucose was measured using a Precision Xtra glucometer (Abbott) to measure 4h fasting blood glucose from the lateral tail vein, as described previously[1].

#### *Tail Bleed*

A tail bleed experiment was performed to test the coagulation ability in tumor bearing mice as previously described[2]. Briefly, tail veins of anesthetized mice were transected and placed in tubes containing warm saline to allow the bleeding time to be visualized and recorded. The weight of the tubes was recorded before and after bleeding to calculate the volume of blood lost.

#### *Coagulation and blood chemistry panels*

Mice were euthanized by isoflurane overdose and blood was collected via cardiac puncture. For coagulation measures, blood was mixed at a blood: sodium citrate ratio (9:1) with 3.2 % sodium citrate. Plasma was then snap frozen after centrifugation at 1500 x g for 15 minutes. The resulting plasma samples were analyzed using an STA Satellite automated coagulation analyzer (Stago). For blood chemistry panel analysis, blood samples were allowed to clot at room temperature for 1 h. The blood was then centrifuged for 10 minutes at 1300 x *g* at 4°C. Resultant serum samples were then analyzed on a Roche Cobas Integra 400 Plus clinical chemistry analyzer.

#### *Gut Permeability Assay*

Mice were fasted overnight and dosed p.o. with 0.15 mL of 80 mg/mL FITC dextran (4kDa). After 4 hours blood was collected via cardiac puncture. Samples were then centrifuged at 5,000 rpm for 10 minutes. Plasma was then diluted in 1x PBS (1:10) and transferred to a black, opaque-bottom 96-well plate. Fluorescence was read on a BioTek Synergy HTX plate reader at ex: 485/em: 530 nm.

#### *Fecal Occult Blood Test*

Fecal matter and blood sample positive control samples analyzed in a luminescent assay for detection of occult bleeding as described previously[3]. Briefly, samples were homogenized with 1 mL of 1X RBC Lysis Buffer (eBioscience) and centrifuged at 8000 x g for 2 minutes at room temperature. A 6% (w/v) solution of Luminol Blend for Forensic Detection of Blood (Innovating Science™ IS5040) was prepared in Milli-Q water. Using an opaque 96-well white plate, 5 µL of each sample were plated in triplicate. To each well, 100 µL of luminol solution was added before luminescence was read in a Bio-Tek Synergy plate reader. Luminescence values were background subtracted and expressed as fold-increase vs. negative control (luminol only).

### Supplementary Figures

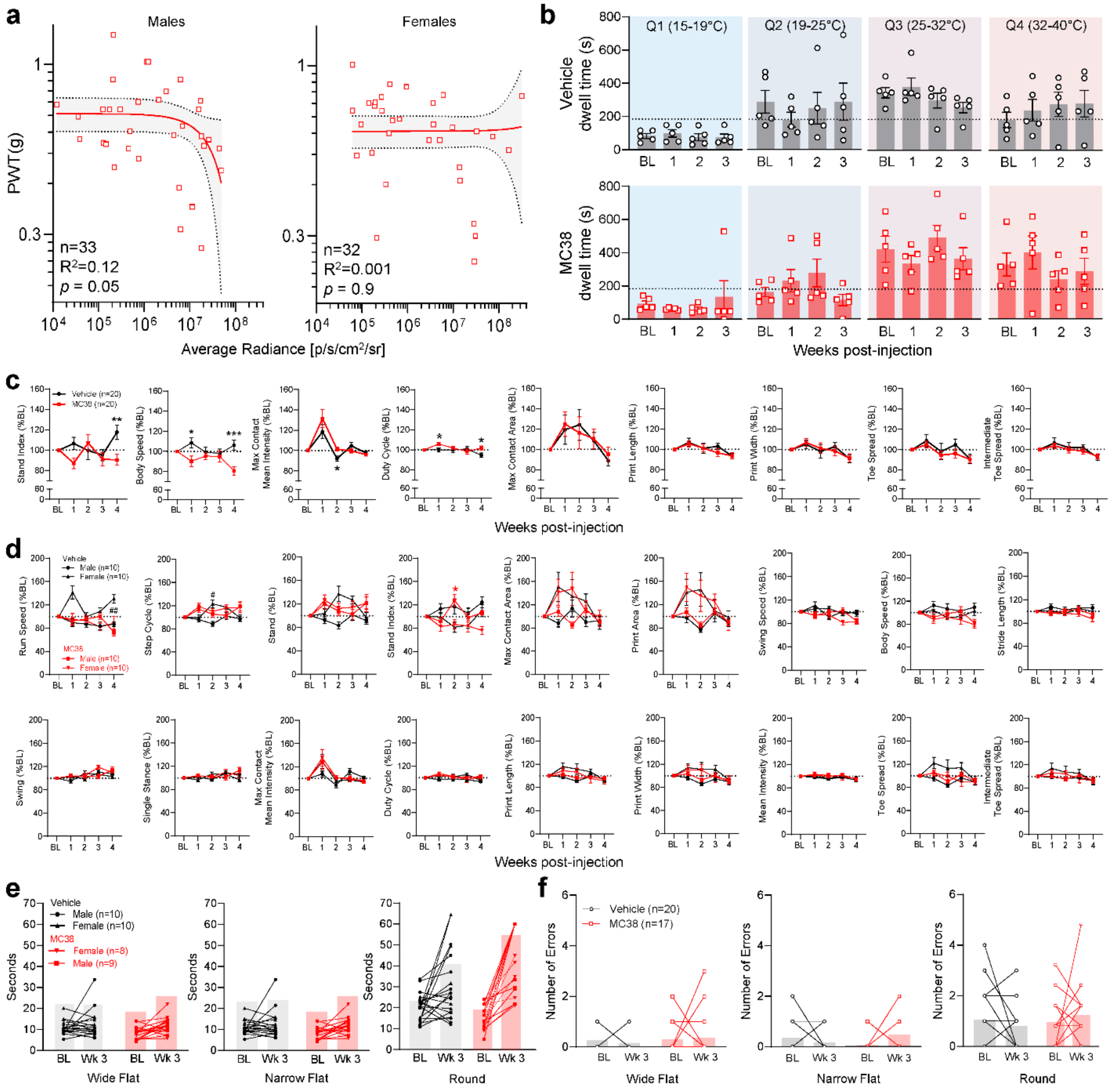


**Supplementary Figure 1. Effect of MC38 tumor growth on sensory sensitivity, gait, and motor coordination.** (**a**) No significant correlation between mean paw withdrawal threshold

(PWT) value and tumor luminescence intensity in male or female MC38 mice. Trendline = linear regression; shaded area = 95% confidence interval. (**b**) Vehicle control mice (top row, grey) display a consistent aversion to the coolest quadrant of the Thermal Gradient Ring (Q1; 15-19°C), with the strongest, most consistent preference for Q3 (25-32°C). MC38 mice (bottom row, red) exhibit a broadly similar temperature preference pattern. Dotted line represents predicted dwell time in the absence of preference (i.e. 750/4 = 187.5 seconds). Error bars = SEM. (**c**) Statistically significant differences in stand index (speed of propulsion phase), body speed (distance traveled in the time taken per step cycle) and mean hindpaw intensity at maximum contact between vehicle-injected and MC38 tumor-bearing mice. No statistically significant differences were observed in duty cycle (proportion of a step cycle spent in stand phase versus swing), hindpaw area at maximum contact, print length/width, or distance between the $1^{st}$ and $5^{th}$ digits ('toe spread') or $2^{nd}$ and $4^{th}$ digits ('intermediate toe spread'). Error bars = SEM, */**/*** $p$ = <0.05 / 0.01 / 0.001, MC38 vs vehicle. Two-way ANOVA, Tukey's multiple comparisons test. (**d**) Significant gait differences between male and female vehicle controls were seen in run speed and step cycle time. Differences between male and female MC38 mice were observed in stand index. All other metrics did not reach statistical significance when comparing sex or injection type. Error bars = SEM, * $p$ = <0.05, MC38 male vs female. # $p$ = <0.05, vehicle male vs female. Two-way ANOVA, Tukey's multiple comparisons test. (**e**) Male and female MC38 mice show similar deterioration in beam walk performance. (**f**) Number of errors (paw slips, inversions, falls) in the beam walk task. Absolute error number increases in the narrow flat and round beams versus the wide flat beam in vehicle controls and MC38 mice.

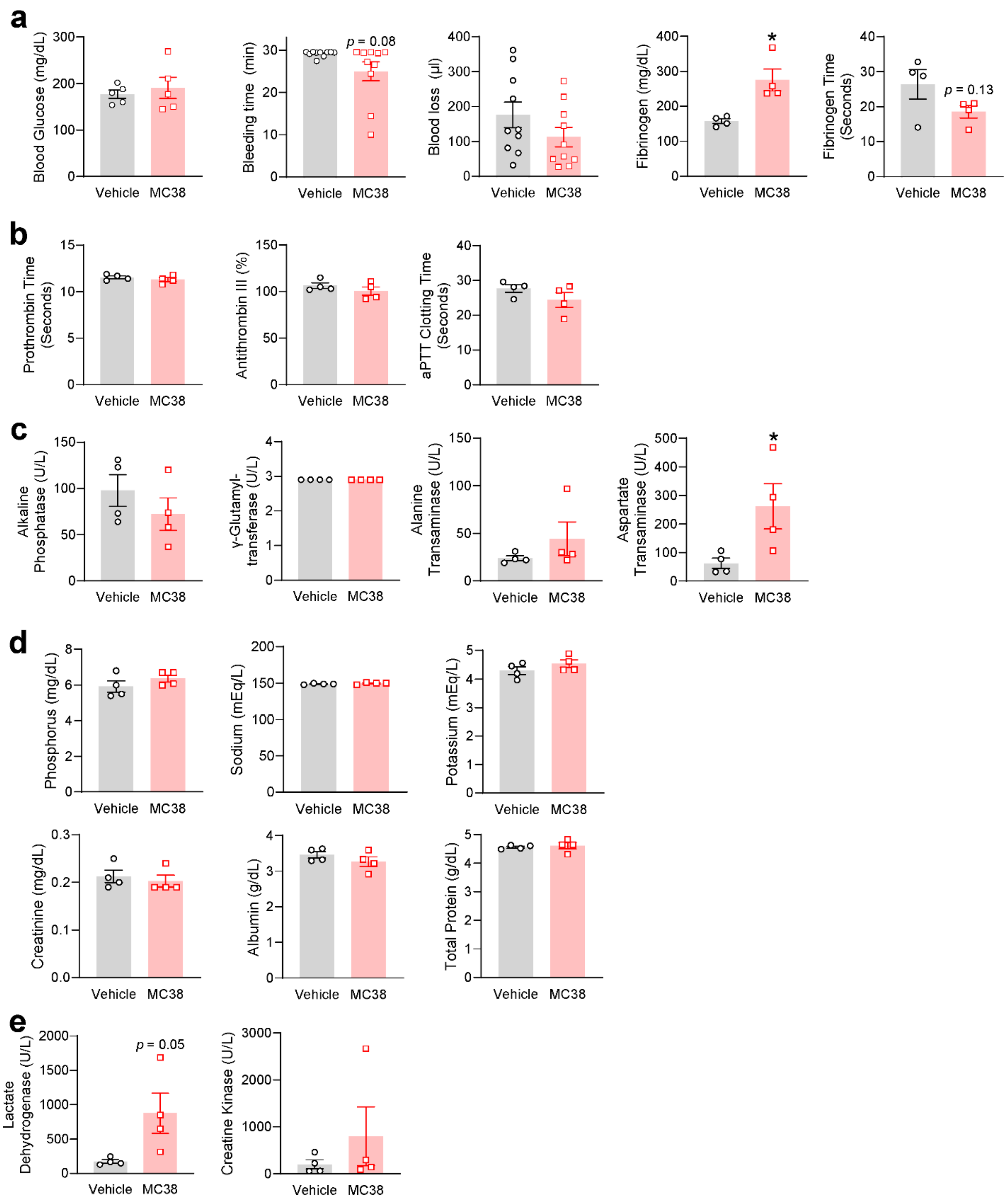


**Supplementary Figure 2. Hematological measures of metabolism, hemostasis, and hepatic/renal health.** (**a**) Three weeks after MC38 injection, blood glucose levels do not differ from vehicle-injected controls. The tail bleed assay shows a strong trend toward reduced bleeding time in MC38 mice, which is associated with a trend toward reduced overall blood loss, significantly elevated fibrinogen, and a trend toward reduced fibrinogen time. (**b**) Prothrombin, antithrombin, and activated partial thromboplastin time (aPTT) do not significantly differ. (**c**)

Liver markers alkaline phosphatase, gamma-glutamyltransferase and alanine transaminase are not significantly elevated in MC38 mice, whereas aspartate transaminase is significantly elevated. (**d**) Renal function as assessed by an electrolyte panel and creatinine appear normal in MC38 mice. Dual liver/kidney function markers albumin and total protein also do not differ. (**e**) A strong trend in upregulation of tissue damage markers lactate dehydrogenase and creatine kinase in MC38 tumor mice. Error bars = SEM. * *p* = <0.05, MC38 versus vehicle, unpaired T-test.

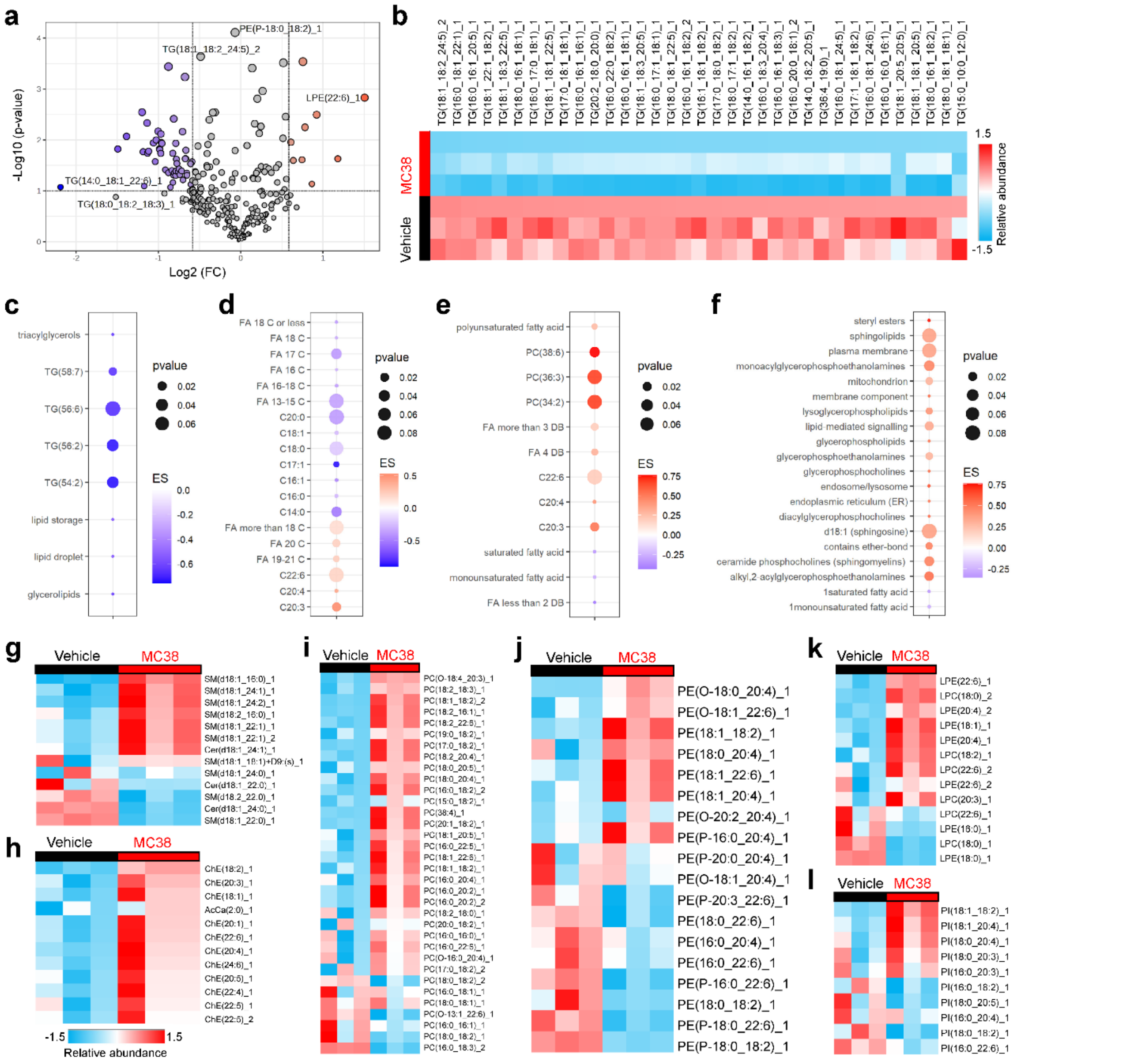


**Supplementary Figure 3. Lipidomic profiling of additional control and MC38 tumor-bearing mouse plasma.** (**a**) Volcano plot displays log2 fold-change and p-value of lipid abundance in MC38 tumor-bearing mouse plasma versus control. (**b**) Heatmap and enrichment score (ES; **c**) shows widespread depletion of triglycerides in tumor-bearing mice. (**d**) Chain length analysis shows an enrichment of long-chain fatty acids and depletion of shorter-chain molecules. (**e**) Saturation index shows enrichment of several polyunsaturated fatty acids alongside depletion of saturated and monounsaturated fatty acids. (**f-g**) Analysis of lipid class enrichment reveals increased ceramides and sphingomyelins. (**h**) Cholesterol esters are enriched in plasma from tumor-bearing mice (**i**) Phosphatidylcholines (PC) are majority enriched in MC38 plasma (**j**) Phosphatidylethanolamines (PE) are majority reduced in MC38 plasma. (**k**) Lysophosphatidylcholine/ethanolamine show similar enrichment/depletion dynamics to PC/PE (**l**) Phosphatidylinositol (PI) abundance is highly variable, with some enrichment and some depletion in MC38 tumor-bearing mice.

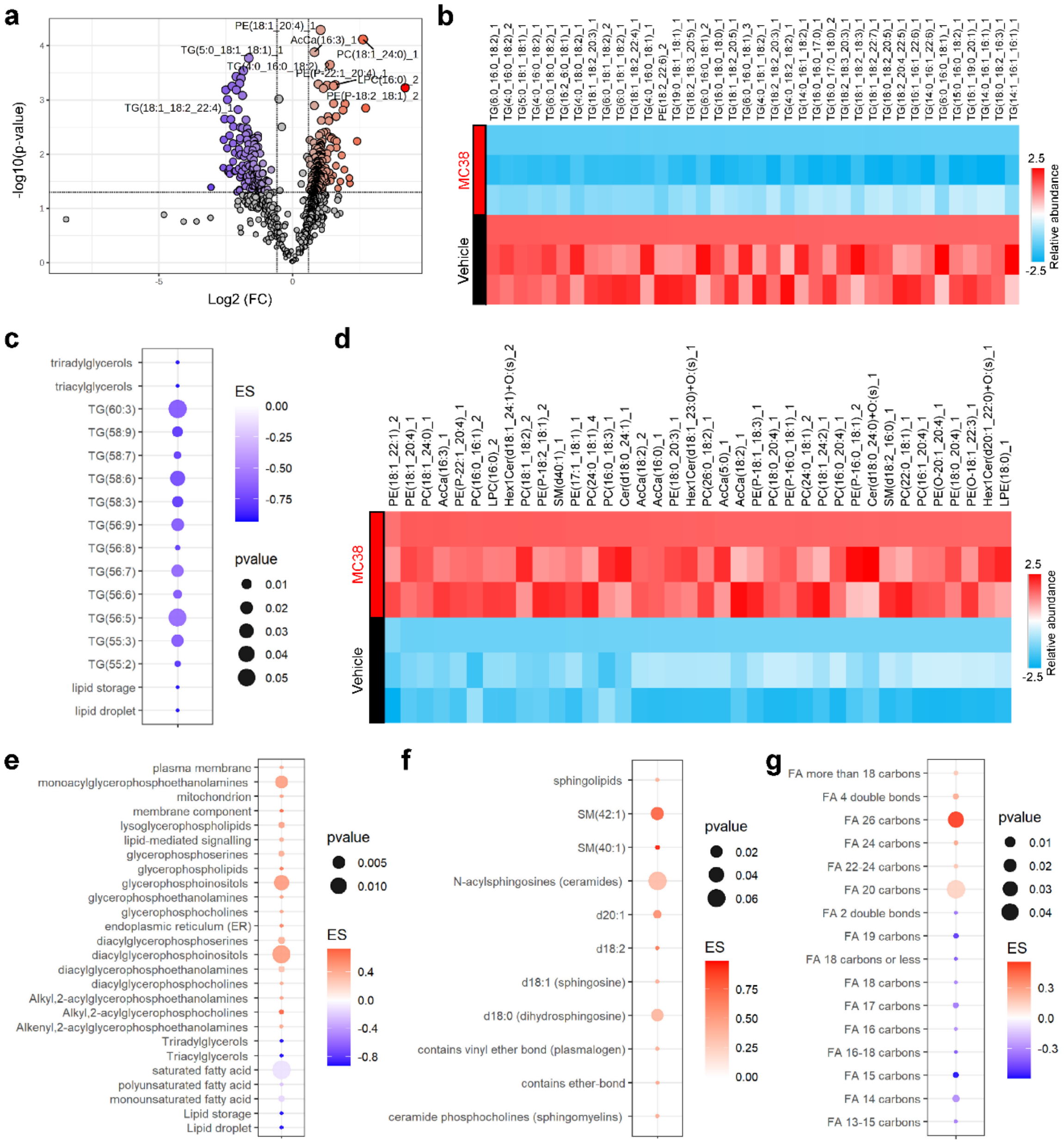


**Supplementary Figure 4. Lipidomic profiling of additional control and MC38 tumor-bearing mouse sciatic nerves.** (**a**) Volcano plot displaying log2 fold-change and p-value of lipid abundance in MC38 tumor-bearing mouse sciatic nerve versus control. (**b-c**) Heatmap and enrichment scores show widespread depletion of triglycerides in tumor-bearing mice. (**d**)

Phosphatidylethanolamines (PE), sphingomyelin, PC, ceramides and hexosylceramides are enriched in sciatic nerves from MC38 tumor mice. (**e**) Analysis of lipid class enrichment reveals increased diacylglycerolipids and reduced triacylglycerols. (**f**) Sphingolipids and ceramides are significantly enriched in MC38 plasma. (**g**) Chain length analysis show enrichment of fatty acids more than 18 carbons in length, and depletion of fatty acids of shorter chain lengths.

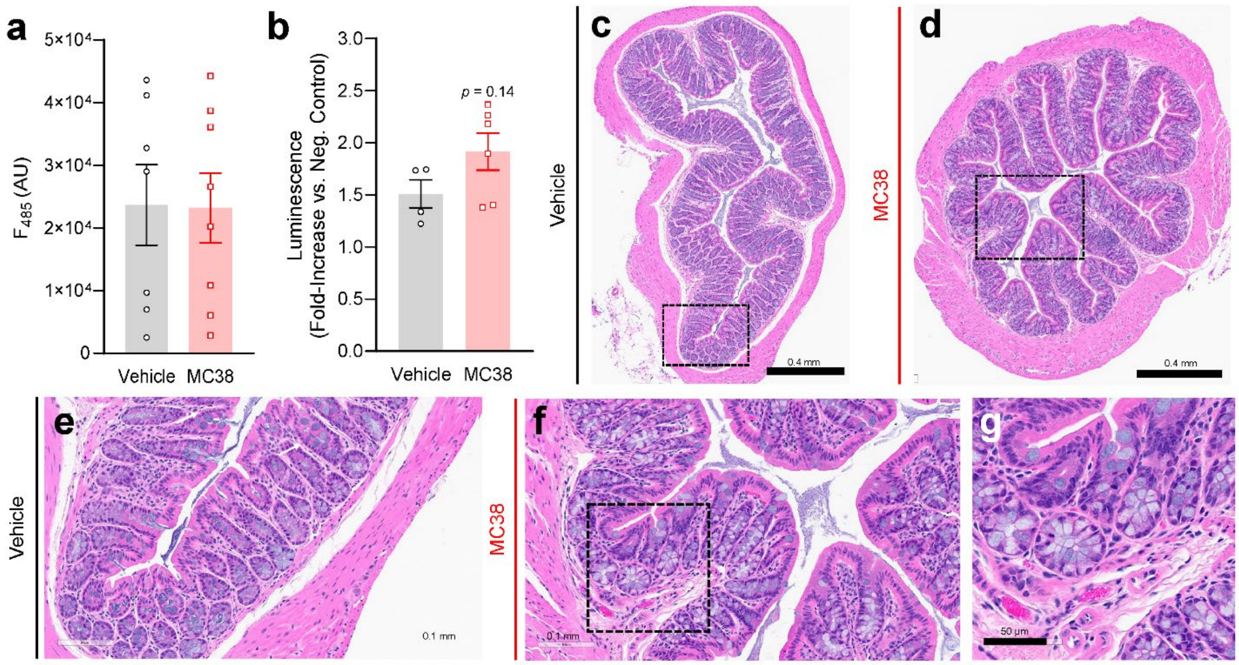


**Supplementary Figure 5. MC38 tumor growth does not increase gut permeability.** (**a**) Oral dosing with a FITC-dextran conjugate was carried out three weeks after MC38 or vehicle injection. Four hours later, blood samples were collected and FITC fluorescence was measured. No significant difference was seen between vehicle and MC38-injected mice. (**b**) Addition of luminol to fecal slurries did not detect any significant increase in hemoglobin content in MC38 tumor mouse feces at three weeks post-injection. (**c-d**) H&E sections of colon appear grossly normal in vehicle and MC38-injected mice. (**e-g**) at higher magnifications an increase in cellularity was detected in MC38 tumor-bearing mice. Magnification 10x (**c-d**), 20x (**e-f**), 40x (**g**). Dotted lines in (**c-d**) show the areas magnified in (**e-f**). Dotted lines in (**f**) show the area magnified in (**g**).

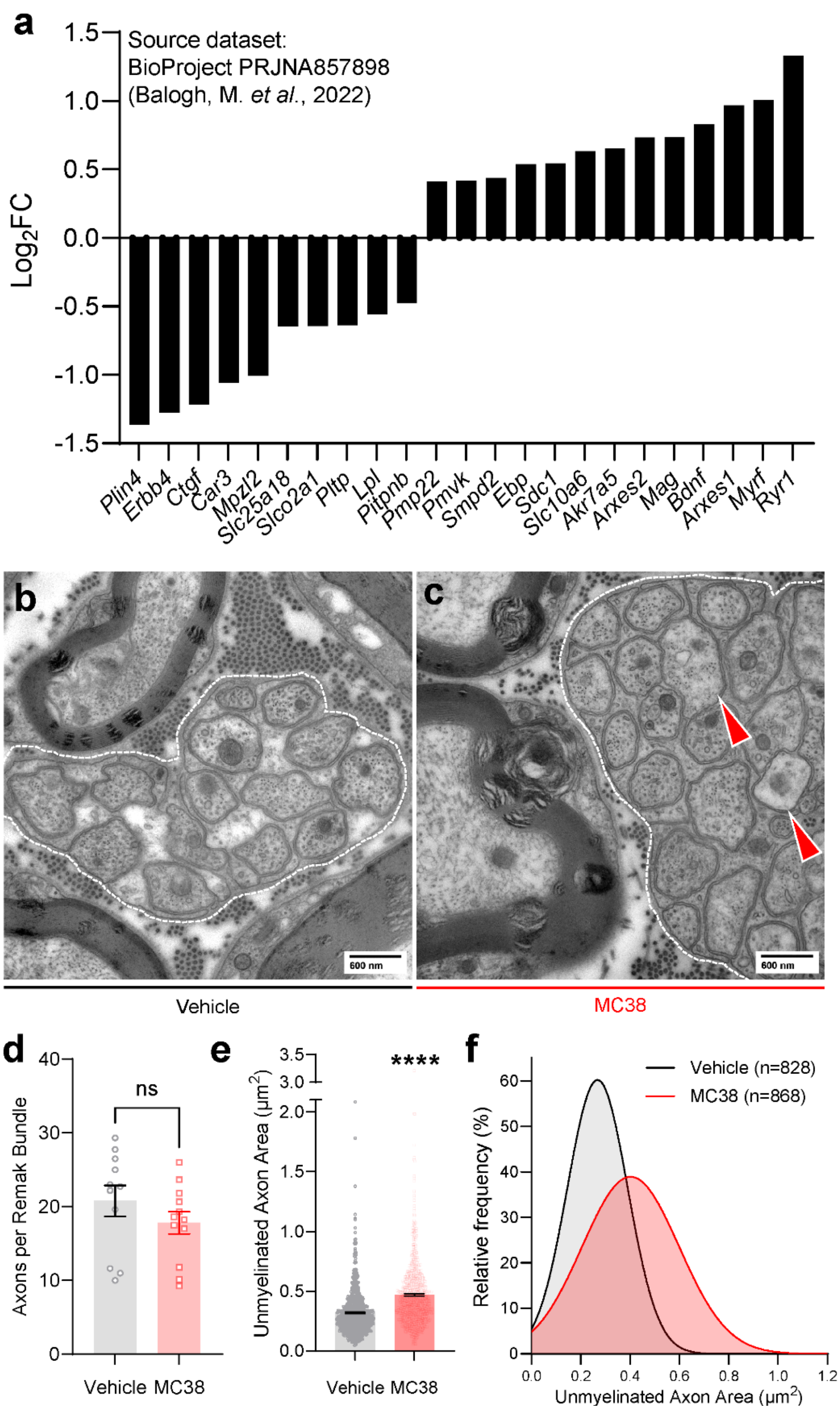


**Supplementary Figure 6. Transcriptional and ultrastructural irregularities associated with MC38 tumor growth.** (**a**) Retrospective analysis of differentially expressed genes was carried out on our previously published dataset (Balogh et al., 2022, BioProject ID: PRJNA857898). Of 559 differentially-expressed genes, 23 were identified as relevant to lipid metabolism and/or

Schwann cell function: *Plin4* (perilipin-4)*, Erbb4* (Erb-B2 receptor tyrosine kinase 4)*, Ctgf* (connective tissue growth factor)*, Car3* (carbonic anhydrase 3)*, Mpzl2* (myelin protein zero-like 2)*, Slc25a18* (solute carrier family 25, member 18)*, Slco2a1* (solute carrier organic anion transporter family member 2a1)*, Pltp* (phospholipid transfer protein)*, Lpl* (lipoprotein lipase)*, Pitpnb* (phosphatidylinositol transfer protein beta)*, Pmp22* (peripheral myelin protein 22)*, Pmvk* (phosphomevalonate kinase)*, Smpd2* (sphingomyelin phosphodiesterase 2)*, Ebp* (emopamil binding protein)*, Sdc1* syndecan 1)*, Slc10a6* (solute carrier family 10, member 6)*, Akr7a5* (aldo-keto reductase family 7 member A5)*, Arxes2* (adipocyte-related X-chromosome expressed sequence 2)*, Mag* (myelin-associated glycoprotein)*, Bdnf* (brain-derived neuroatrophic factor)*, Arxes1* (adipocyte-related X-chromosome expressed sequence 1)*, Myrf* (myelin regulatory factor) and *Ryr1* (ryanodine receptor 1). (**b**) Sciatic nerve TEM shows small-diameter, unmyelinated axons organized into Remak bundles (**b**), with some axons in MC38 samples showing signs of swelling (red arrowheads, **c**). The number of axons per Remak bundle does not differ significantly in MC38 mice (**d**), but there is a significant increase in mean cross-sectional area of unmyelinated axons (**e**), also detectable as a rightward shift in the frequency distribution (**f**). Error bars = SEM. **** $p$ = <0.0001, MC38 versus vehicle, unpaired T-test.

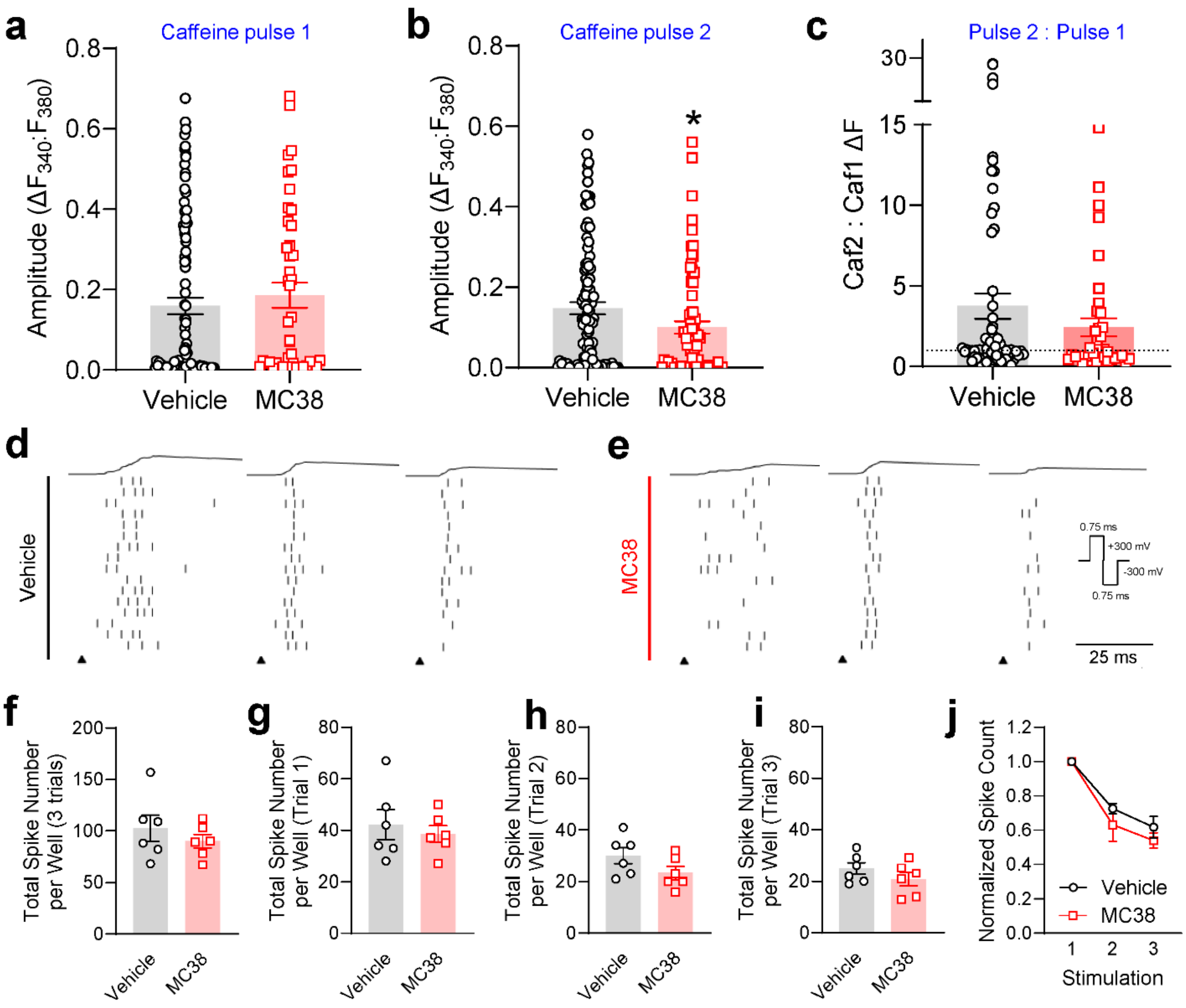


**Supplementary Figure 7. Amplitude of caffeine-induced $Ca^{2+}$ responses and electrically-evoked activity in sensory neurons from control and MC38 tumor mice.** (**a**) Caffeine pulse 1 (pre-KCl depolarization) evokes $Ca^{2+}$ flux to a similar extent in vehicle and MC38-treated mice. (**b**) Caffeine pulse 2 (post-KCl depolarization) evokes a smaller average peak $Ca^{2+}$ influx in MC38-treated mice. (**c**) The ratio of pulse 2 to pulse 1 is above 1 in vehicle and MC38-treated mice, suggesting that KCl depolarization fills ER $Ca^{2+}$ stores beyond their resting state in both conditions. However, the ratio shows a trend toward lower values in MC38-treated mice. Error bars = SEM, * $p$ = <0.05, unpaired T-test. (**d**) Representative raster plots of one well of cultured DRG neuron responses to three bouts of electrical stimulation. The indicated waveform was applied simultaneously via four of the sixteen electrodes in each well (carats). After removal of stimulation artifacts (< 2ms), this stimulation evoked 1-4 spikes per unit across all electrodes within ~25ms. Similar activity was evoked from DRG cultures isolated from vehicle controls (**e**) and MC38 mice. (**f**) Total number of spikes evoked per well across all three trials did not significantly differ, nor was there a greater degree of 'run-down' in evoked activity in MC38 neuron cultures (**g-j**).

**Supplementary Files:**

**Supplementary Video Files 1-3: Male vehicle vs MC38 tumor, week 3, wide flat, narrow flat, round beams.**

Representative video recordings of vehicle control and MC38 mice (males) traversing the easy (wide flat), intermediate (narrow flat) and difficult (round) beams.

**Supplementary Video Files 4-6: Female vehicle vs MC38 tumor, week 3, wide flat, narrow flat, round beams.**

Representative video recordings of vehicle control and MC38 mice (females) traversing the easy (wide flat), intermediate (narrow flat) and difficult (round) beams.

**Supplementary Table 1: Plasma Lipidomic Analysis**

Fold-change and adjusted p-values of all significantly increased and decreased lipid species in MC38 mouse plasma (3 weeks) versus control plasma.

**Supplementary Table 2: Sciatic Nerve Lipidomic Analysis**

Fold-change and adjusted p-values of all significantly increased and decreased lipid species in MC38 mouse sciatic nerve (3 weeks) versus control sciatic nerve.

**Supplementary Table 3: Sciatic Nerve Proteome Profile Analysis**

Fold-change of all detected factors in lysates of MC38 (3 weeks) versus control sciatic nerves.
